# SPACE: Spatially variable gene clustering adjusting for cell type effect for improved spatial domain detection

**DOI:** 10.1101/2024.08.23.609477

**Authors:** Sikta Das Adhikari, Nina G. Steele, Brian Theisen, Jianrong Wang, Yuehua Cui

**Affiliations:** Department of Computational Mathematics, Science and Engineering, Michigan State University, East Lansing, MI; Department of Statistics and Probability, Michigan State University, East Lansing, MI; Department of Surgery, Henry Ford Pancreatic Cancer Center, Henry Ford Hospital, Detroit, MI; Department of Pathology, Wayne State University, Detroit, MI; Department of Oncology, Wayne State University, Detroit, MI; Department of Pharmacology and Toxicology, Michigan State University, East Lansing, MI; Department of Pathology, Henry Ford Health, Detroit, MI; Department of Statistics and Probability, Michigan State University, East Lansing, MI, 48824, USA

**Keywords:** Spatial Transcriptomics, Spatially variable genes, Spatial domain, Gene spatial pattern classification

## Abstract

Recent advances in spatial transcriptomics have significantly deepened our understanding of biology. A primary focus has been identifying spatially variable genes (SVGs) which are crucial for downstream tasks like spatial domain detection. Traditional methods often use all or a set number of top SVGs for this purpose. However, in diverse datasets with many SVGs, this approach may not ensure accurate results. Instead, grouping SVGs by expression patterns and using all SVG groups in downstream analysis can improve accuracy. Furthermore, classifying SVGs in this manner is akin to identifying cell type marker genes, offering valuable biological insights. The challenge lies in accurately categorizing SVGs into relevant clusters, aggravated by the absence of prior knowledge regarding the number and spectrum of spatial gene patterns. Addressing this challenge, we propose SPACE, SPatially variable gene clustering Adjusting for Cell type Effect, a framework that classifies SVGs based on their spatial patterns by adjusting for confounding effects caused by shared cell types, to improve spatial domain detection. This method does not require prior knowledge of gene cluster numbers, spatial patterns, or cell type information. Our comprehensive simulations and real data analyses demonstrate that SPACE is an efficient and promising tool for spatial transcriptomics analysis.

**Key Points:**

- SPACE eliminates the need for prior knowledge about the number of gene clusters, known cell types, or the quantity of SVGs to identify clusters for downstream analysis.
- SPACE offers a method to effectively leverage SVGs for low-dimensional embedding within each cluster to improve the accuracy of spatial domain detection.
- The efficiency and utility of the SPACE algorithm have been validated across multiple datasets and simulations, demonstrating its effectiveness in producing meaningful and interpretable results.

## 1 Introduction

Recent advancements in spatially-resolved transcriptomics (SRT) technology have revolutionized our ability to acquire comprehensive gene expression data for thousands of genes across tissue locations in multiple samples. The number of genes and spatial resolution vary depending on the specific technology employed. However, regardless of the technology and resolution, spatial transcriptomic data facilitate the exploration of various biological questions.

Often a fundamental initial step in the analysis of SRT data involves identifying spatially variable genes (SVGs). These genes exhibit expression level variations either across the entire tissue or within predefined spatial domains. In recent years, there has been an abundance of research and the development of new methods to address the challenge of detecting SVGs[1][2]. Although the detection of SVGs lets us visualize the spatial patterns in the tissue which might offer some level of biological insights about the tissue of interest, the main use of SVGs lies in downstream analysis, specifically for spatial domain detection. Spatial domains are distinct and functionally specialized anatomical structures within tissue, each distinguished by unique local characteristics including cell-type composition, transcriptome heterogeneity, and cell-cell interactions[3][4][5]. Detecting these domains is crucial for understanding their collaborative role in tissue functions and development stages. To achieve this, a set number of top SVGs is typically selected, and spatial domains are identified using these top SVGs.

However, using an arbitrary number of top SVGs might not represent all the spatial patterns exhibited by the SVGs. As previously argued [6], Some dominant patterns may overshadow less pronounced yet relevant patterns. Previous methods, like SpatialDE[7], SPARK [8] and Sepal[6], attempted to classify SVGs into groups with similar spatial patterns, aiding in a more holistic representation of results. It’s important to highlight that classifying SVGs is a challenging task requiring specialized methods. Simple clustering approaches are inadequate as they overlook spatial information [7]. However, in existing methods challenges arise regarding the selection of the unknown number of spatial pattern groups and selection of other parameters, as well as the unclear impact of classification on downstream analysis. Here, we propose an efficient method SPACE to classify SVGs into clusters and explore the benefits of this clustering step in the final goal of spatial domain detection.

The concept of clustering SVGs carries a significant biological rationale. Researchers have already identified many SVGs as the markers for specific cell types [9],[10],[11],[12], yet detecting these cell type-specific SVGs presents a formidable challenge. As distinct spatial patterns are associated with distinct cell type compositions, it follows that different cell type-specific SVGs would manifest distinct spatial patterns. Consequently, clustering SVGs with similar patterns can be viewed as a means of segregating distinct cell type-specific SVGs.

SPACE is a Gaussian process-based method that initially identifies SVGs and then establishes a dependency map among these SVGs using an intuitive approach (see Materials and Methods 2). This map links each SVG with other SVGs exhibiting similar spatial patterns, ultimately clustering similarly expressed SVGs together (see Figure 1A). The resulting SVG-clusters from the SPACE algorithm can serve as inputs for further downstream analysis.

**Figure 1:**
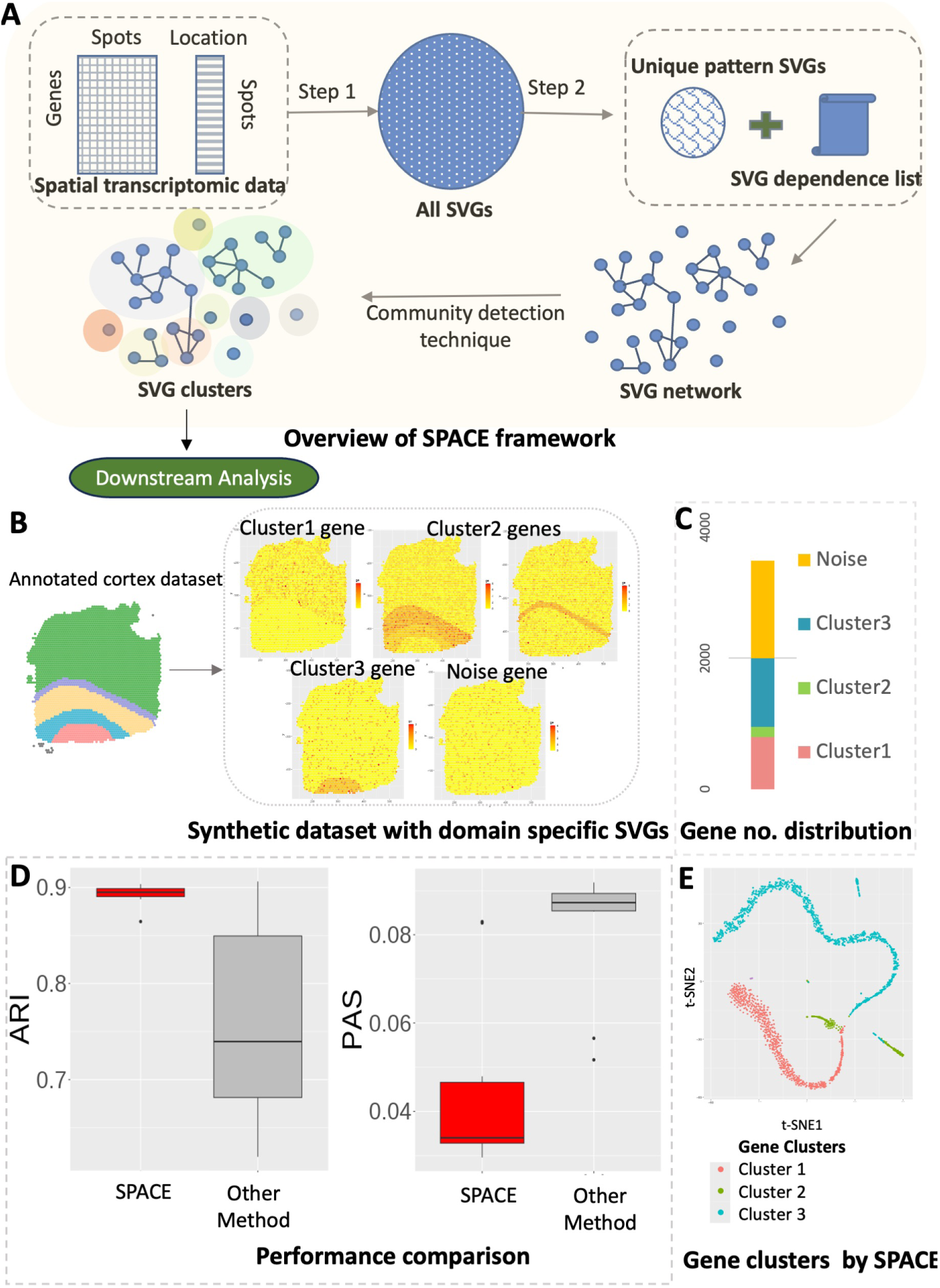
(A) Schematic overview of the SPACE framework. (B-E) An example simulation based on a synthetic dataset: (B) The synthetic dataset is generated based on a real dataset with annotated spatial domains. Three types of domain-specific SVGs and noise genes are created. (C) The distribution of the number of each type of gene is presented. (D) Evaluation based on ARI (higher is better) and PAS score (lower is better). (E) Visualization by the t-SNE plot: shown is for a randomly selected simulation result. The gene clusters identified by SPACE are highlighted with distinct colors. The plot illustrates that genes within the clusters are densely grouped together and separated from genes belonging to other clusters.

For performing downstream analysis, we employ a well-known dimension reduction technique tailored for spatial data, SpatialPCA[13], to derive low-dimensional embeddings specific to each SVG-cluster. These embeddings are subsequently utilized for spatial domain detection. In each example considered in this study, whether through simulation setups or real data analysis, we compare our findings with those obtained from the SpatialPCA framework, which generates low-dimensional embeddings for all top SVGs collectively and subsequently performs the same spatial domain detection step. This comparison aims to elucidate the advantages of the gene clustering step facilitated by SPACE.

To evaluate the performance of SPACE, we conducted a comparison using a synthetic dataset derived from real human DLPFC cortex data, with known annotations of its 5 layers (4 prefrontal cortex layers and the white matter). See Supplementary Figure 2 for more details on the synthetic data generation. Figure 1B illustrates the domain-based SVG clusters within the synthetic dataset: cluster 1 represents genes predominantly expressed in layer 1, cluster 2 includes genes from cortex layers 2, 3, or 4, while cluster 3 comprises genes overexpressed in the white matter region. The distribution of the number of genes across clusters is uneven in this scenario as shown in figure 1C, as frequently evident in real datasets. Upon repeating the analysis on 10 simulated synthetic datasets, and considering ARI scores (Adjusted Rand Index, higher the better) and PAS scores (Percentage of abnormal spots, lower the better, see Materials and Methods 2) (see Figure 1D), we observe that our framework provides better domain detection results compared to the SpatialPCA framework. Additionally, Figure 1E displays a t-SNE plot [14] representing genes from a randomly chosen simulation outcome. The genes are color-coded according to the gene cluster label identified by SPACE. This visualization demonstrates that genes within each cluster are packed together, indicating the accuracy of gene classification by SPACE.

In this paper, we conducted two types of simulation studies: firstly, to assess the accuracy of the SPACE framework for SVG classification, and secondly, to evaluate the accuracy of domain detection based on the detected SVG clusters by SPACE using synthetic datasets mimicking real-world scenarios. In addition, we analyzed three publicly available datasets and one newly acquired pancreatic ductal adenocarcinoma (PDAC) SRT dataset. The publicly available datasets include: 1)The DLPFC human cortex annotated dataset[15], comprising 12 samples; 2)The HER2 human breast tumor annotated dataset[16](used one sample); and 3)The dataset from the study of human breast cancer biopsies[17]. The newly acquired dataset is the pancreatic ductal adenocarcinoma(PDAC) dataset, which comes with rudimentary annotation. The application of SPACE and the detection of domains aligns well with the rough annotation. Overall, findings from simulation studies and real data analyses across multiple datasets affirm that utilizing SVG clusters and, consequently, the SPACE framework can markedly enhance the performance of spatial domain detection. This approach holds significant promise to unveil hidden spatial patterns that provide novel insights into the spatial heterogeneity of tissue samples.

## 2 Materials and Methods

In a typical spatial transcriptomics setup, the dataset comprises gene expression measures or counts for *m* genes distributed across *N* known spatial coordinates or spots. Suppose *y* = (*y*_1_*, y*_2_*, …, y_N_* ) represents the gene expression profiles or counts for a particular gene across spatial coordinates (referred to as samples or spots), denoted by *s* = (*s*_1_*, …, s_N_* ). The spatial locations are typically represented as two-dimensional coordinates, i.e., *s_i_* = (*s_i_*_1_*, s_i_*_2_), although coordinates of any dimensionality can be employed. The primary objective of spatially variable gene (SVG) detection models is to identify which genes, among the *m* genes, exhibit spatial variability across the tissue. In essence, the key goal is to determine whether the gene expression measure *y* is dependent on or related to the spatial locations where the gene expression measures are sampled.

Consider a scenario where there are *k_d_* spatial domains within the tissue of interest. A spatial domain represents a distinct region or area within the tissue characterized by unique molecular signatures or gene expression patterns. These patterns may arise from specific factors such as cellular composition, anatomical organization, spatial arrangement, or functional attributes.

In reality, the specific spatial domains are typically unknown. However, each domain may be defined with a set of SVGs exhibiting characteristic gene expression patterns in proximity to these domains. To accurately reconstruct the underlying domain structure, it is beneficial to group SVGs based on their spatial patterns and utilize all SVG groups in downstream analysis, rather than relying solely on an arbitrary subset of all SVGs.

With this objective in mind, we present the SPACE framework, organized into two primary steps. The first step involves the selection of SVGs, while the second step utilizes the SVGs selected in the first step to generate an SVG dependency list and subsequently create clusters of SVGs based on this information (see 1A). It is worth noting that there exists a plethora of SVG detection techniques in the literature [1] [2], and any of these methods can substitute for the first step in the framework. However, methods that rigorously control the False Discovery Rate (FDR) are preferable, as they enhance accuracy in the subsequent stages of analysis.

### 2.1 Step 1: Selecting SVGs

Like the majority of the SVG detection methods this step is based on the Gaussian process (GP) regression model which models the normalized gene expression *y* for a given gene assuming the following multivariate normal model:

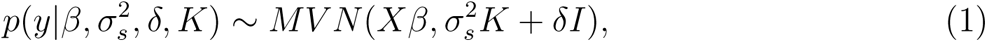

where the covariance term is decomposed into a spatial and a non-spatial part, with *δI* and 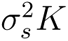 representing the non-spatial and spatial covariance matrix, respectively. The (*i, j*)*^th^* element in the kernel matrix *K* denotes the spatial similarity between the *i^th^* and *j^th^* spot calculated based on the corresponding coordinates *s_i_* and *s_j_*. The choice of the kernel function plays a very important role in detecting the spatial correlation presented in the gene expressions. *X^N×k^* represents the covariate matrix, while *β^k×^*^1^ denotes the array of corresponding coefficients. This model can incorporate up to *k −* 1 covariates, such as cell type information or domain structure information. However, often such information is either unavailable or deemed untrustworthy. Hence, in practice, we typically employ *X* solely as the intercept.

When evaluating the existence of spatial patterns within the data, an assessment is made by testing the alternative hypothesis, which suggests the presence of a spatial variance in the model, against the null hypothesis, where the spatial variance component is zero, indicating the absence of spatial variability. This comparison between the model fitted under the null and alternative hypotheses forms the basis of a significance testing procedure. This often involves conducting significance tests and drawing conclusions based on p-values in frequentist approaches. In model (1), testing if a gene is spatially variable is equivalent to testing 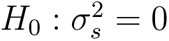.

Within this framework, we use a straightforward score test to test the underlying hypothesis, and a p-value is calculated. More information regarding the test is provided in the supplementary material. Prior studies [8, 18] indicate that Gaussian and Cosine kernels are adept at capturing a wide spectrum of distinct spatial gene expression patterns. Hence, we utilize 10 different kernels (5 Cosine and 5 Gaussian kernels with varying parameter values) following the approach established by SPARK [8]. Denote the number of detected SVGs by *m*_1_ in step 1.

### 2.2 Step 2: Classifying SVGs by spatial patterns

This stage categorizes the SVGs identified in step 1 into cohesive groups. While conventional clustering algorithms can segregate genes into distinct groups based on gene expression, they invariably disregard spatial information. Hence, we require more sophisticated algorithms to classify spatially variable genes more precisely [7].

Intuitively, if two spatially variable genes *y*_1_ and *y*_2_ exhibit similar spatial patterns, they should be correlated which is equivalent to assume *y*_1_ = *α*_0_ + *α*_1_*y*_2_ + *ɛ*, where 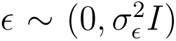 is a random noise term with arbitrary variance 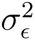 and some constants *α*_0_*, α*_1_. Hence, while testing gene *y*_1_, if we use gene *y*_2_ as a covariate in model (1) and test the same alternative hypothesis described in step 1, we would expect gene *y*1 to be less or even not significant depending on how strong the correlation between *y*_1_ and *y*_2_ is. On the other hand, if after using another gene *y*_3_ as a covariate in the model we still find gene *y*_1_ to be significant, that would imply that gene *y*_1_ and gene *y*_3_ have different spatial patterns. Given that many SVGs are cell type marker genes [9][10][11][12], we would expect gene *y*_1_ and *y*_2_ belong to the same cell type while gene *y*_3_ belongs to a different cell type. However, our algorithm does not require such knowledge (often not available in spot-level SRT data) as detailed below.

With this intuition, for each SVG *j, j* = 1*, · · ·, m*_1_, we can select a list of genes *S_j_* which are correlated to gene *j* and use them as covariates in the model 1. If no of genes in *S_j_* exceeds 3, we choose the top *k_j_* principal components and include them as covariates in the model. Typically, *k_j_* is chosen such that at least 80% of the total variance is explained by the top *k_j_* principal components. With this, model (1) becomes:

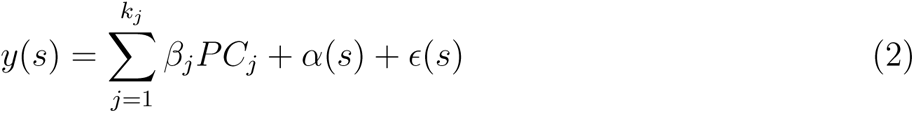

where *PC_j_*represents the *j^th^*principal component, 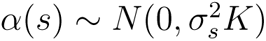 and *ɛ*(*s*) *∼ N* (0*, δI*).

There might be different ways of selecting related genes. It could be done by choosing a threshold based on a correlation measure, either linear or nonlinear, such as Pearson correlation, Spearman correlation, distance correlation or kernel correlation. Alternatively, one can run a penalized regression with LASSO[19], Elastic net [20][21] or MCP penalty[22] or perform sure independence screening[23] to select genes correlated with gene *j*.

In real applications, many SVGs represent marker genes for cell types, exhibiting spatial expression patterns that reflect the distribution of various cell types. Given that marker genes for a given cell type often exhibit similar or strongly correlated gene expression distributions, employing this model adeptly controls for the cell-type specific effect, a factor typically challenging to ascertain or quantify in real-world settings as the cell type information is typically unknown in spot-level SRT data.

Employing this model on all the genes, we can get a set of genes with unique patterns (showing significance under model (2)) and a gene dependency list. From the list, we come up with a weighted graph structure, where nodes are SVGs and two nodes are connected if they are correlated. By applying a clustering algorithm such as leiden [24], groups of genes are determined and the unique genes (which are not part of any gene group) create singleton sets. The full algorithm steps are provided in the supplementary file.

### 2.3 Downstream analysis: spatial domain detection

Spatially resolved transcriptomics serve a crucial role in identifying tissue or region sub-structures through domain detection analysis. Numerous frameworks have been developed for this purpose, to name a few, SpaGCN[25], SpatialPCA [13] and BayesSpace[26]. In our analysis, we opted for SpatialPCA[13] due to its proven superiority in performance over other available algorithms. Furthermore, to conduct domain detection post identification of SVG clusters using our framework, we require an effective low-dimensional representation of the dataset, a task efficiently facilitated by SpatialPCA. SpatialPCA effectively extracts a low-dimensional representation of spatial transcriptomic data while preserving both biological signals and spatial correlation structures. This condensed representation can serve as an input for efficient clustering algorithms, such as the Louvain [27] or Walktrap algorithm [28], facilitating the clustering of spots and thereby identifying spatial domains. The steps for obtaining spatial domains in the SpatialPCA workflow include: 1) Select the top 3000 SVGs and calculate spatial PCs based on these genes, and 2) Use the top 20-30 spatial PCs for spatial clustering using algorithms like Louvein or walktrap.

In our approach, we utilize our framework to identify SVG groups with similar spatial patterns. Within each SVG group, we compute the spatial PCs and aggregate the top spatial PCs from each group to form the final embedding. These aggregated spatial PCs are then used as input in the clustering algorithm to detect spatial domains. The differences between our method and the existing ones are that existing methods use low-dimensional embeddings based on all top SVGs, while our method gets low-dimensional embeddings within each cluster. The within-cluster embeddings can capture unique spatial structures and hence lead to improved spatial domain detection.

### 2.4 Measuring accuracy of domain detection

Detecting domains essentially involves assigning a cluster label to each of the *N* spots in the tissue sample. Once a framework is implemented for detecting spatial domains, it becomes crucial to measure its accuracy against the ground truth domain labels. We primarily employ a standard clustering evaluation metric, the Adjusted Rand Index (ARI), to assess the similarity between the predicted domain labels and the true labels. Additionally, we utilize the PAS (Percentage of Abnormal Spots) score to quantify the clustering performance of spatial domain detection, following the approach outlined in [13]. This score gauges the randomness of spots located outside their clustered spatial domain computed as the proportion of spots with a cluster label differing from at least six of their ten neighboring spots. A lower PAS score reflects greater homogeneity within spatial clusters.

## 3 Results

### 3.1 Simulation Study

We conducted two simulation studies: one to demonstrate the effectiveness of the SVG clustering performance and another to assess whether the SVG clusters identified by SPACE are beneficial for enhancing domain detection accuracy.

#### 3.1.1 Evaluation of the clustering performance

To illustrate SPACE’s capability to accurately classify SVGs, we devise the simulation scenario I in which 100 normalized gene expression datasets were simulated, each consisting of 53 genes (labeled g1-g53) and 2000 spots following model 1 with no covariates. The first 10 genes (g1-g10) represent independent noise genes, not correlated with any other gene in the dataset. The next set of 10 genes (g11-g20) exhibit strong correlations among themselves but do not show any spatial pattern. The third (g21-g30), fourth (g31-g40), and fifth (g41-g50) sets of genes represent spatially variable genes with three distinct spatial patterns (Spatial Pattern 1, Spatial Pattern 2, and Spatial Pattern 3, respectively). Figure 2A exhibits the six sets of representative genes with distinct spatial patterns. Genes are correlated within the spatial pattern 1-3 as exhibited in Figure 2B, and the spatial effect strength increases with the gene index. For example, g21 and g30 are correlated and share the same spatial pattern, but the spatial effect is stronger in g30 compared to g21. The correlation between the genes within a gene group could have different structures. They could display Compound Symmetry (CS) if any pair of genes within a group have the same correlation, i.e., *ρ_ij_* = *ρ*. Alternatively, they could demonstrate a first-order Autoregressive (AR(1)) pattern if the correlation between two genes decays as their distance increases, i.e., *ρ_ij_* = *ρ^|i−j|^*. Figure 2B exhibits CS correlation structure. The last three genes, g51, g52, and g53, each exhibit a unique spatial pattern (Spatial Pattern 4, Spatial Pattern 5, and Spatial Pattern 6, respectively). As the spatial pattern strength increases within each spatial gene group (pattern1-pattern3), the SVG detection power converges towards 1, as expected (see Figure 2C). Furthermore, upon comparing this outcome with the performance of the most efficient comparable SVG detection method, nnSVG[29] (where both nnSVG and SPACE use normalized gene expressions), we observed that SPACE detects spatial genes slightly more effectively (see Supplementary Figure S4). The SPACE framework not only demonstrates efficacy in identifying the true SVGs (g21-g53) but also adeptly categorizes them. The Adjusted Rand Index (ARI) scores, calculated for each simulated dataset, are computed based on the cluster labels of detected SVGs and their true cluster labels - cluster near 1 in the violin plot in Figure 2D, indicating high accuracy of gene clustering. The false discovery rates were also monitored in this study. Here false discovery happens when any of the non-spatial genes (g1-g20) appears in any of the final gene clusters. FDR is calculated as the number of false discoveries divided by the total number of SVG discoveries. As indicated in Figure 2E, the FDR values are predominantly distributed near 0, indicating high accuracy of the results.

**Figure 2:**
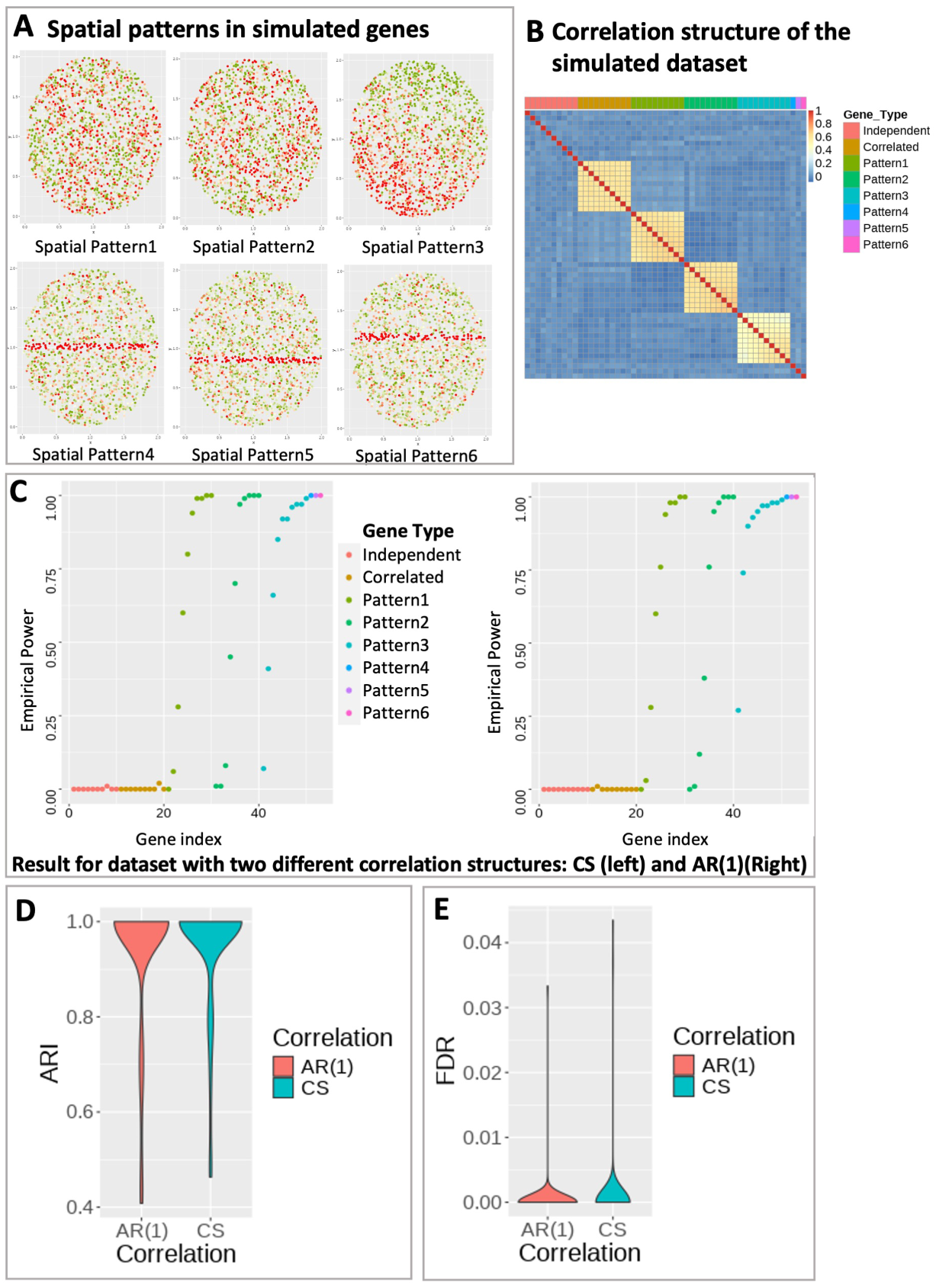
Simulation setting for SPACE. A) Six representative genes each with a distinct spatial pattern in the simulated dataset. Green and Red color represents low and high gene expression. B) Correlation heatmap of the simulated dataset with compound symmetry correlation structure within gene groups. Independent: uncorrelated gene group for genes without any spatial pattern; Correlated: correlated gene group for genes without any spatial pattern. Pattern 1-3: correlated gene group for genes with spatial pattern 1-3. Pattern 4-6: single gene with spatial pattern 4-6. C) Empirical power of the SVG detection step at detecting SVGs for the simulated datasets with compound symmetry (Left) and AR(1) (Right) correlation structure within each gene group. D) The distribution of ARI values based on predicted gene clusters for simulation under each correlation structure. E) Empirical FDR distribution for simulation under each correlation structure. Here, false discovery occurs when Correlated or Independent genes show up in any of the final gene clusters.

#### 3.1.2 Evaluation of the spatial domain detection performance

To showcase how the outputs of the SPACE algorithm aid in the downstream analysis of spatial domain detection and enhance its accuracy, we generated synthetic datasets based on the annotated human DLPFC data with sample ID 151670. Supplementary Figure S2 details the steps involved in generating the synthetic data, ensuring that key features of the original dataset are preserved, such as the distribution of means and variances of all genes. The dataset is annotated with 5 layers (4 prefrontal cortex layers and the white matter layer). After filtering out sparse genes, we ended up with 4,865 genes whose expressions were measured in 3484 spots. We first randomly selected 2,000 genes which were converted to SVGs in the generated dataset (See supplementary Figure 2). The rest of the genes were converted to random noise genes with no specific pattern. Among the 2,000 SVGs, three distinct spatial domain structures were represented (see 1B and 1C): 800 SVGs correspond to the first cluster, wherein genes are predominantly expressed in the cortex layer 1; 150 SVGs exhibit overexpression in cortex layers 2, 3, or 4; and the remaining 1,050 SVGs from cluster 3 predominantly display expression in the white matter domain region. In such 10 simulated synthetic datasets, we applied our framework, SPACE, to identify gene clusters which were further leveraged for domain detection. We compared the domain detection results of SPACE with the default one by SpatialPCA without an additional clustering step. As we mentioned previously in the introduction 1, our framework significantly improves the spatial domain detection accuracy, as evidenced by ARI scores and PAS scores (see 1D. Additionally, t-SNE plot[14] from a randomly chosen simulation result for the genes displays the accuracy of the gene clustering performance of SPACE. The t-SNE plots for all 10 simulation results were given in Supplementary Figure S3.

### 3.2 Real data analysis

#### 3.2.1 Human DLPFC 10x Genomics Visium dataset

We applied the SPACE algorithm to the human dorsolateral prefrontal cortex (DLPFC) data[15] generated by Visium from 10x Genomics. Publicly available datasets from 12 human DLPFC tissue samples, obtained from three individuals, can be accessed and downloaded from the link: http://spatial.libd.org/spatialLIBD/. We directly downloaded the processed datasets from the SpatialPCA Github repository, available at: https://github.com/shangll123/SpatialPCA_analysis_codes. These samples, on average, encompassed 3,973 spots, each manually annotated to one of the six prefrontal cortex layers or white matter. We illustrated the method focusing on two samples with ID 151670 and 151673, which contain expression measurements of 33,538 genes across 3,498 spots and 33,538 genes across 36,39 spots, respectively. We also analysed the other 10 samples and the results are available in the supplementary files.

We applied our framework to detect the spatial domains (see methods 2) for each of the samples. We also followed the SpatialPCA framework provided in https://github.com/shangll123/SpatialPCA_analysis_codes without separating genes into clusters to detect spatial domains to compare with our results. The comparison is based on the ARI scores, utilizing the provided annotations for each sample. We also compared the PAS scores for all the samples. In Figure 3A, sample 151670 is presented with annotated spatial domains as the ground truth (left), spatial domains detected by SpatialPCA (middle), and spatial domains detected by our framework (right). The ARI values for the detected domains by SpatialPCA and our framework (0.34 and 0.69, respectively, as shown in Figure 3E), along with the PAS scores (0.013722 and 0.006289, respectively, depicted in Figure 3F), highlight the superior performance of our framework over SpatialPCA.

**Figure 3:**
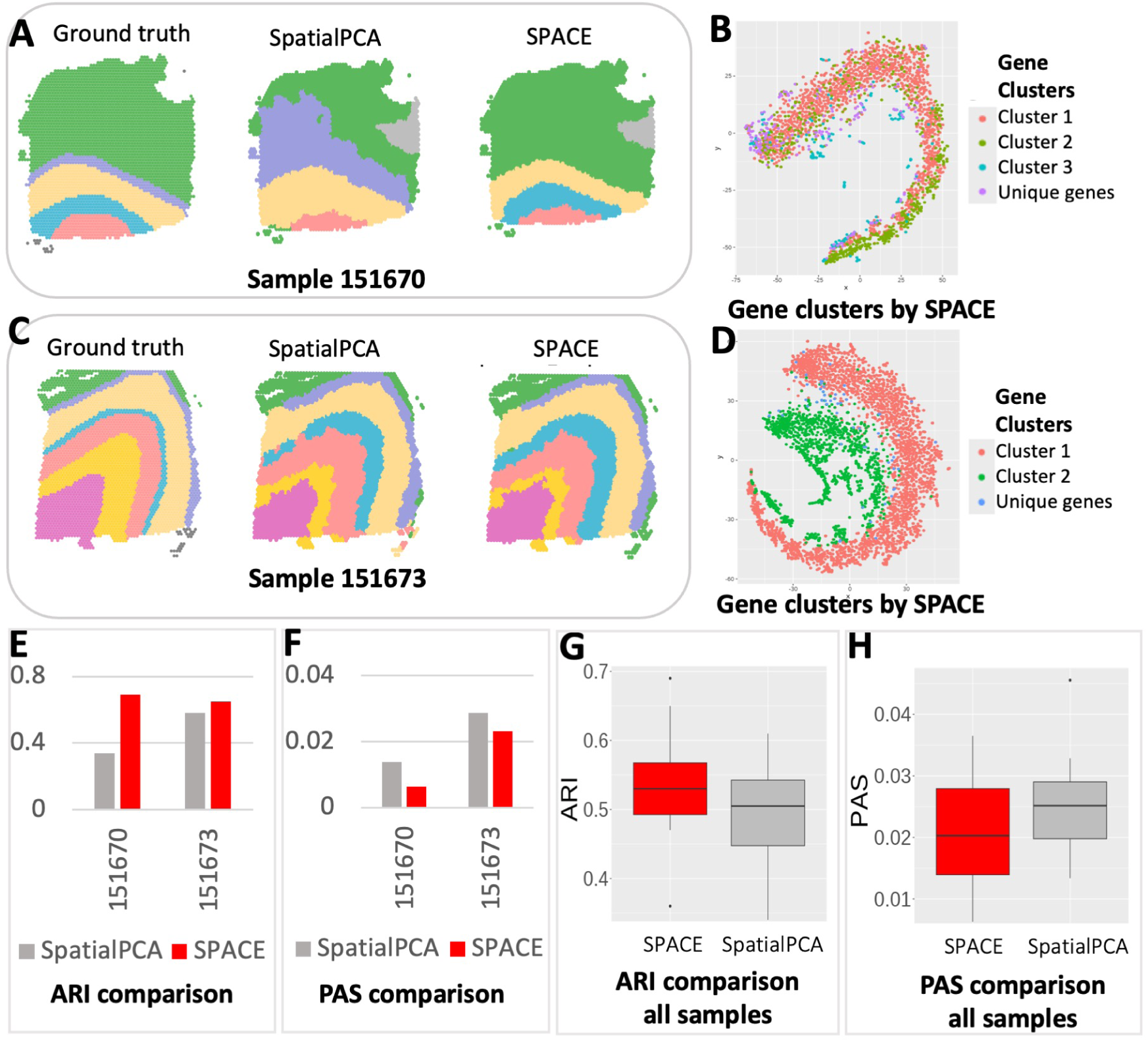
Spatial domain detection analysis results of the Human cortex data from DLPFC: (A) The detected spatial domains for Sample 151670: Annotated domain used as the ground truth (left), by SpatialPCA (middle), and by our framework (Right). (B) The t-SNE plot for all the SVGs detected by SPACE for Sample 151670. SVGs are colored based on cluster labels calculated by SPACE. (C) The spatial domains detected for Sample 151673: Annotated domain used as the ground truth (left), by SpatialPCA (middle), and by our framework (Right). (D) The t-SNE plot for all the SVGs detected by SPACE for Sample 151673. The fact that SVGs with similar colors are close to each other but are away from those with different colors indicates accurate gene clustering. (E) Comparison of ARI score (higher the better) between SpatialPCA and our framework based on these two samples. (F) Comparison of PAS score (lower the better) between SpatialPCA and our framework based on these two samples. (G) Comparison of ARI score between SpatialPCA and our framework based on all 12 samples. (H) Comparison of PAS score between SpatialPCA and our framework based on all 12 samples.

Similarly, in Figure 3C, the spatial domains for sample 151673 are displayed in the same order: annotated domains, spatial domains by SpatialPCA, and spatial domains by our framework. The ARIs for SpatialPCA and our framework are 0.58 and 0.65, respectively and the PAS scores are 0.028579 and 0.023083, respectively. Combining the results from all the samples, our framework significantly enhances spatial domain detection compared to SpatialPCA (see Figure 3G, 3H) as confirmed by the increase in the median ARI score.

We visualized the SVG clusters identified by SPACE through the t-SNE plots in Figure 3B and 3D for the two samples. These visualizations highlight the resemblance among genes within the same cluster and the disparity between genes from separate clusters, emphasizing the efficacy of SPACE in delineating SVG clusters.

#### 3.2.2 Analysis of HER2 breast tumor data

We applied SPACE to another dataset, the HER2-positive breast tumor data[16], initially comprising 36 tumor datasets from eight individuals (patients A-H), each consisting of 3 or 6 sections. Following SpatialPCA analysis, we selected the H1 dataset, encompassing 15,030 genes measured across 613 spatial locations. We utilized the pre-processed dataset available in the SpatialPCA repository, containing 10,053 genes across 607 spots, and omitted datasets from other samples due to sparse gene expressions or minimal spot coverage. SPACE identified 268 SVGs grouped into three main clusters, along with a few unique pattern genes. The three main gene clusters consist of 84, 48, and 117 genes respectively. The t-SNE plot of the SVGs (see Figure 4B) exhibits a similar pattern for genes within the same cluster in close proximity to each other. Comparing spatial domains detected by SpatialPCA and SPACE based on the ARI value, our framework shows better performance (ARI=0.48) than SpatialPCA (ARI=0.44), thereby improving domain detection accuracy (see 4C).

**Figure 4:**
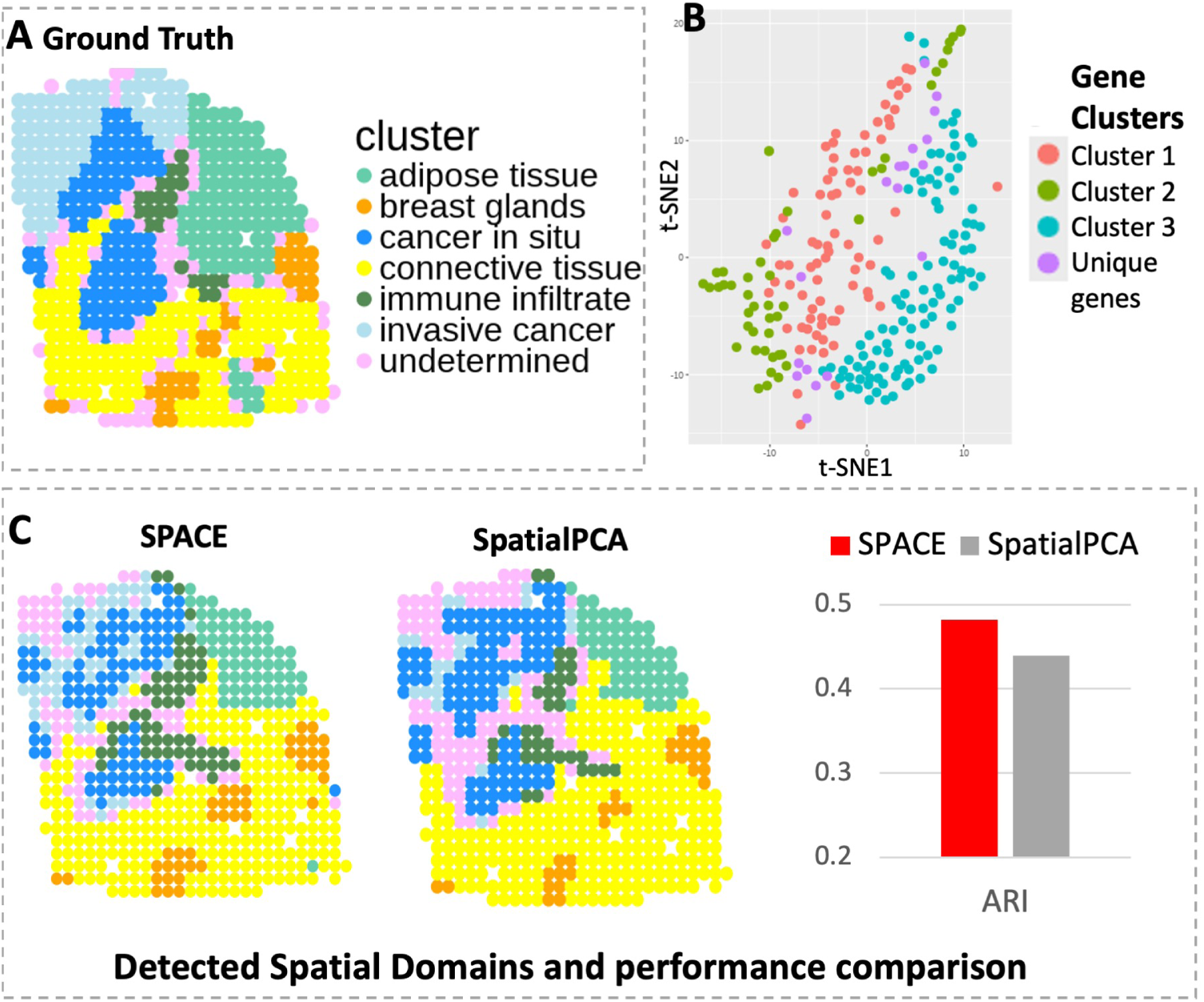
Analysis of the HER2 data. (A) Annotated spatial layers considered as ground truth, showcasing 6 known tissue components including cancer-related spots in blue shades. (B) The t-SNE plot illustrating SVGs detected by SPACE, with distinct colors representing distinct SVG clusters. (C) Spatial domains detected by SPACE and SpatialPCA, with respective ARI scores of 0.48 and 0.44.

#### 3.2.3 Analysis of the PDAC data

Our final analysis was conducted on a Pancreatic ductal adenocarcinoma (PDAC) dataset, obtained from the Henry Ford Health System^1^. This dataset comprises gene expression measurements for 17,943 genes across 3,142 spots. Following standard filtering and normalization procedures, we applied our SPACE framework to identify SVG clusters and detect spatial domains. Three primary SVG clusters were identified, each showcasing differential expressions in distinct tissue regions. Representative genes from these clusters are depicted in supplementary Figure S8.

For each SVG cluster, we extracted low-dimensional embeddings (top SpatialPCs from SpatialPCA), combined them, and applied the Leiden algorithm to identify spatial domains. We repeated this process for the SpatialPCA framework, utilizing the top 20 Spatial PCs from the top 3,000 SVGs. Due to the absence of spot annotations, the exact number of spatial domains is unknown. Therefore, we employed the same Leiden algorithm with a resolution parameter set at the default value of 1 for the SpatialPCA framework, rather than using algorithms typically utilized by SpatialPCA that require a predetermined number of domains.

The dataset includes a rough annotation (see 5A) highlighting the tumor (marked in red) and non-tumor (marked in yellow) regions of interest. Figures 5B and 5C display the predicted domains by SpatialPCA and SPACE frameworks, respectively. As the precise annotation labels for each spot are unavailable, we cannot compute scores like ARI to compare spatial domain detection accuracy between the two methods. However, through visual inspection, the domains detected by SPACE appear more accurate, effectively capturing the most important regions. Tumor-containing regions identified by SPACE are notably smaller and more accurate compared to those identified by SpatialPCA.

**Figure 5:**
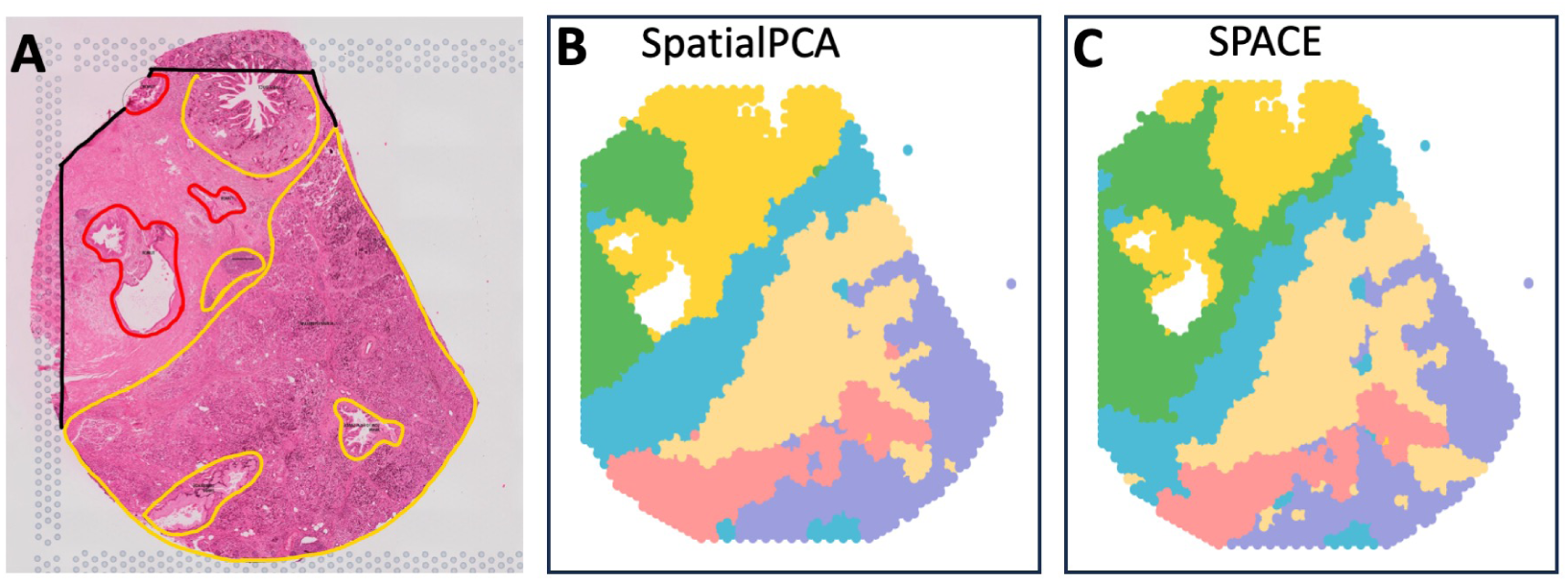
Analysis of the PDAC data: (A) Rough annotation of the tissue depicting different important regions. Tumor regions are highlighted in red, while non-tumor yet significant regions are marked in yellow. (B) Spatial domains detected by SpatialPCA, utilizing the calculation of spatial PCs and employing the Leiden algorithm for clustering without presupposing the number of clusters. (C) Spatial domain detection by SPACE, involving the aggregation of SVG-cluster-specific spatial PCs and utilizing the Leiden algorithm for clustering without presupposing the number of clusters.

The SPACE algorithm identified three primary clusters of spatially variable genes (comprising 3931, 2751, and 1781 genes, respectively) in the PDAC dataset, each providing significant biological insights. Supplementary Figure S8 demonstrates that genes in Cluster 1 are predominantly overexpressed mostly in the tumor adjacent regions, while genes in Clusters 2 and 3 are overexpressed around tumor regions. Consequently, pathway enrichment analysis shown in Figure 6 reveals that Cluster 1 included pathways highly suggestive of changes in the metabolic state of the cells. Changes in the metabolic state of tumor and tumor adjacent regions is a well-established feature in PDAC [30],[31]. Cluster 3 is uniquely enriched for inflammatory pathways specific to T cell signaling, a feature not observed in Cluster 1. These data suggest there is a specific spatial distribution of immune (and T and NK cells specifically) within tumor adjacent regions, in agreement with previous work [32]. Immune suppression is a key aspect of PDAC [33], and our pipeline may lead to new understandings of the mechanisms driving this aspect of the disease in the future. Cluster 2 displayed a mixed result with pathways identified both on inflammatory signaling, and cell to cell communication (i.e. tight junctions). As PDAC is highly desmoplastic, this result may reflect the abundant cancer associated fibroblasts (CAFs) present in this cancer type that signal to tumor cells and promote or restrict tumor growth depending on the subtype of CAF present.

**Figure 6:**
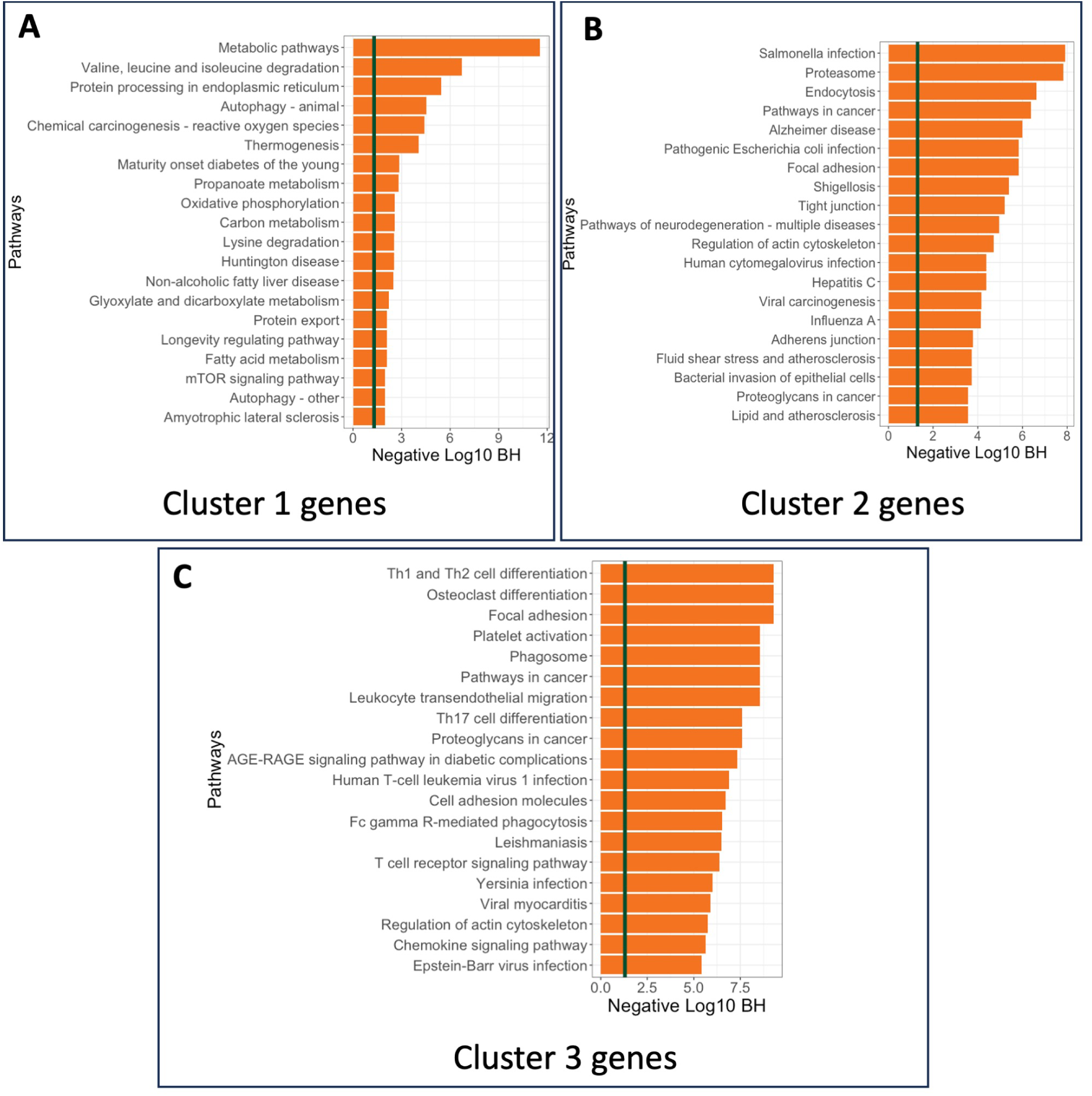
Enrichment analysis of the PDAC data: KEGG Pathway enrichment analysis of genes from A) Cluster 1, B) Cluster 2, and C)Cluster 3 are showcased.

On the other hand, a similar enrichment analysis on the top 3000 SVGs(see Supplementary Figure S9(A) or the group of all SVGs(see Supplementary Figure S9(B) shows that the gene sets are enriched in a mixture of different pathways, which does not clearly indicate the functional dependency between spatial gene patterns and their biological mechanisms. The unique enrichment of cluster 3 SVGs in Figure 6C for inflammatory pathways specific to T cell signaling would be impossible to discern if all SVGs or the top 3000 SVGs were selected. This demonstrates why using SVG clusters instead of just the top genes provides more biological insights and suggests that using SVG clusters might be a better option for downstream analysis, carrying more biological information.

## 4 Discussion

In recent years, the exploration and analysis of spatial data have reached unprecedented heights, offering diverse insights into biological systems. Central to this endeavor is the identification of SVGs, which serve as pivotal components in understanding tissue organization and function. However, merely detecting SVGs does not inherently yield substantial biological insights. Rather, their significance lies in their dual role: 1) SVGs are used for spatial domain detection and 2) Some SVGs serve as markers for specific cell types. Traditional approaches often struggle to achieve precise spatial domain detection, and discerning cell type-specific SVGs amidst the data noise poses a formidable task. Our proposed framework, SPACE, addresses these challenges to improve spatial domain detection without requiring cell type information.

By implementing SPACE, we effectively detect SVG clusters that can be interpreted as clusters of cell-type SVGs. Remarkably, this identification is accomplished without the need for complex cell-type deconvolution techniques, streamlining the analysis process while providing biologically meaningful insights. Through extensive real data analysis and simulation studies, we have demonstrated the efficacy of SPACE in enhancing spatial domain detection accuracy, validating its utility in spatial transcriptomic analysis.

The implementation of SPACE involves two sequential steps, with the initial phase focusing on SVG detection. This step can be performed using any existing SVG detection technique and the alteration can be done without changing any code for SPACE. The next step focuses on finding gene cluster which involves using the Leiden community detection algorithm [24] (details in Supplementary). The detailed step-by-step code and results are provided on our GitHub repository https://github.com/wangjr03/SPACE. For all the results in the paper, the parameters involved in the community detection algorithm were not tuned, and default values were used. However, it has been observed that tuning the parameters can further improve spatial domain detection accuracy in some cases. Nonetheless, tuning the parameters in this instance poses certain challenges. Looking ahead, there are ample opportunities to refine and expand upon the SPACE framework. Future enhancements may involve simplifying and scaling up the SVG detection and cluster detection methods to accommodate larger and more complex datasets. Additionally, continued refinement of the framework will enable researchers to extract deeper insights from spatial transcriptomic data, advancing our understanding of tissue biology and disease mechanisms.

In conclusion, the use of SVG clusters generated by SPACE represents a crucial advancement in spatial transcriptomic analysis. As we continue to refine and evolve this methodology, it is poised to become an indispensable tool for dissecting the spatial complexity of biological systems and unraveling the intricate interplay between genes, cells, and tissues.

## Data Availability

All relevant codes for reproducing each step of the real data analysis and simulation study results are available on our GitHub repository: https://github.com/wangjr03/SPACE. The publicly accessible datasets and their sources are provided in the data folder. Please note that the PDAC data used in this study has not been published and is not publicly available at the time of submission.

## Supplementary Data statement

This manuscript includes supplementary materials.

## Acknowledgement

We thank Mr. Ian Loveless for his contribution to the PDAC sample alignment. We also thank MSU iCER for providing the high-performance computing infrastructure.

## Footnotes

1 Institutional Review Board approval is maintained for 16150 at Henry Ford Hospital for The Translational and Clinical Research Center Biorepository

## References

[1] Sikta Das Adhikari, Jiaxin Yang, Jianrong Wang, and Yuehua Cui. Recent advances in spatially variable gene detection in spatial transcriptomics. Comput. Struct. Biotechnol. J., February 2024.

[2] Zhijian Li, Zain M Patel, Dongyuan Song, Guanao Yan, Jingyi Jessica Li, and Luca Pinello. Benchmarking computational methods to identify spatially variable genes and peaks. bioRxivorg, December 2023.

[3] Charles Swanton. Intratumor heterogeneity: evolution through space and time. Cancer research, 72(19):4875–4882, 2012.

[4] Michalina Janiszewska. The microcosmos of intratumor heterogeneity: the space-time of cancer evolution. Oncogene, 39(10):2031–2039, 2020.

[5] David T Scadden. Nice neighborhood: emerging concepts of the stem cell niche. Cell, 157(1):41–50, 2014.

[6] Alma Andersson and Joakim Lundeberg. sepal: Identifying transcript profiles with spatial patterns by diffusion-based modeling. Bioinformatics, 37(17):2644–2650, 2021.

[7] Valentine Svensson, Sarah A Teichmann, and Oliver Stegle. Spatialde: identification of spatially variable genes. Nature methods, 15(5):343–346, 2018.

[8] Shiquan Sun, Jiaqiang Zhu, and Xiang Zhou. Statistical analysis of spatial expression patterns for spatially resolved transcriptomic studies. Nature methods, 17(2):193–200, 2020.

[9] Miranda V Hunter, Reuben Moncada, Joshua M Weiss, Itai Yanai, and Richard M White. Spatially resolved transcriptomics reveals the architecture of the tumor-microenvironment interface. Nature communications, 12(1):6278, 2021.

[10] Qianghu Wang, Baoli Hu, Xin Hu, Hoon Kim, Massimo Squatrito, Lisa Scarpace, Ana C DeCarvalho, Sali Lyu, Pengping Li, Yan Li, et al. Tumor evolution of glioma-intrinsic gene expression subtypes associates with immunological changes in the microenvironment. Cancer cell, 32(1):42–56, 2017.

[11] Yi Zhang, Guanjue Xiang, Alva Yijia Jiang, Allen Lynch, Zexian Zeng, Chenfei Wang, Wubing Zhang, Jingyu Fan, Jiajinlong Kang, Shengqing Stan Gu, et al. Metatime integrates single-cell gene expression to characterize the meta-components of the tumor immune microenvironment. Nature communications, 14(1):2634, 2023.

[12] Srivatsan Raghavan, Peter S Winter, Andrew W Navia, Hannah L Williams, Alan DenAdel, Kristen E Lowder, Jennyfer Galvez-Reyes, Radha L Kalekar, Nolawit Mulugeta, Kevin S Kapner, et al. Microenvironment drives cell state, plasticity, and drug response in pancreatic cancer. Cell, 184(25):6119–6137, 2021.

[13] Lulu Shang and Xiang Zhou. Spatially aware dimension reduction for spatial transcriptomics. Nature communications, 13(1):7203, 2022.

[14] Laurens Van der Maaten and Geoffrey Hinton. Visualizing data using t-sne. Journal of machine learning research, 9(11), 2008.

[15] Kristen R Maynard, Leonardo Collado-Torres, Lukas M Weber, Cedric Uytingco, Brianna K Barry, Stephen R Williams, Joseph L Catallini, Matthew N Tran, Zachary Besich, Madhavi Tippani, et al. Transcriptome-scale spatial gene expression in the human dorsolateral prefrontal cortex. Nature neuroscience, 24(3):425–436, 2021.

[16] Alma Andersson, Ludvig Larsson, Linnea Stenbeck, Fredrik Salmén, Anna Ehinger, Sunny Z Wu, Ghamdan Al-Eryani, Daniel Roden, Alex Swarbrick, Åke Borg, et al. Spatial deconvolution of her2-positive breast cancer delineates tumor-associated cell type interactions. Nature communications, 12(1):6012, 2021.

[17] Patrik L Ståhl, Fredrik Salmén, Sanja Vickovic, Anna Lundmark, Jośe Ferńandez Navarro, Jens Magnusson, Stefania Giacomello, Michaela Asp, Jakub O Westholm, Mikael Huss, et al. Visualization and analysis of gene expression in tissue sections by spatial transcriptomics. Science, 353(6294):78–82, 2016.

[18] Jiaqiang Zhu, Shiquan Sun, and Xiang Zhou. Spark-x: non-parametric modeling enables scalable and robust detection of spatial expression patterns for large spatial transcriptomic studies. Genome biology, 22(1):184, 2021.

[19] Robert Tibshirani. Regression shrinkage and selection via the lasso. Journal of the Royal Statistical Society Series B: Statistical Methodology, 58(1):267–288, 1996.

[20] Jerome Friedman, Trevor Hastie, and Rob Tibshirani. Regularization paths for generalized linear models via coordinate descent. Journal of statistical software, 33(1):1, 2010.

[21] Noah Simon, Jerome Friedman, Trevor Hastie, and Rob Tibshirani. Regularization paths for cox’s proportional hazards model via coordinate descent. Journal of statistical software, 39(5):1, 2011.

[22] Cun-Hui Zhang. Nearly unbiased variable selection under minimax concave penalty. 2010.

[23] Jianqing Fan and Jinchi Lv. Sure independence screening for ultrahigh dimensional feature space. Journal of the Royal Statistical Society Series B: Statistical Methodology, 70(5):849–911, 2008.

[24] Vincent A Traag, Ludo Waltman, and Nees Jan Van Eck. From louvain to leiden: guaranteeing well-connected communities. Scientific reports, 9(1):5233, 2019.

[25] Jian Hu, Xiangjie Li, Kyle Coleman, Amelia Schroeder, Nan Ma, David J Irwin, Edward B Lee, Russell T Shinohara, and Mingyao Li. Spagcn: Integrating gene expression, spatial location and histology to identify spatial domains and spatially variable genes by graph convolutional network. Nature methods, 18(11):1342–1351, 2021.

[26] Edward Zhao, Matthew R Stone, Xing Ren, Jamie Guenthoer, Kimberly S Smythe, Thomas Pulliam, Stephen R Williams, Cedric R Uytingco, Sarah EB Taylor, Paul Nghiem, et al. Spatial transcriptomics at subspot resolution with bayesspace. Nature biotechnology, 39(11):1375–1384, 2021.

[27] Vincent D Blondel, Jean-Loup Guillaume, Renaud Lambiotte, and Etienne Lefebvre. Fast unfolding of communities in large networks. Journal of statistical mechanics: theory and experiment, 2008(10):P10008, 2008.

[28] Pascal Pons and Matthieu Latapy. Computing communities in large networks using random walks. In Computer and Information Sciences-ISCIS 2005: 20th International Symposium, Istanbul, Turkey*, October 26-28*, *2005. Proceedings 20*, pages 284–293. Springer, 2005.

[29] Lukas M Weber, Arkajyoti Saha, Abhirup Datta, Kasper D Hansen, and Stephanie C Hicks. nnsvg for the scalable identification of spatially variable genes using nearestneighbor gaussian processes. Nature communications, 14(1):4059, 2023.

[30] Zeribe C Nwosu, Matthew H Ward, Peter Sajjakulnukit, Pawan Poudel, Chanthirika Ragulan, Steven Kasperek, Megan Radyk, Damien Sutton, Rosa E Menjivar, Anthony Andren, et al. Uridine-derived ribose fuels glucose-restricted pancreatic cancer. Nature, 618(7963):151–158, 2023.

[31] Christopher J Halbrook and Costas A Lyssiotis. Employing metabolism to improve the diagnosis and treatment of pancreatic cancer. Cancer cell, 31(1):5–19, 2017.

[32] Nina G Steele, Eileen S Carpenter, Samantha B Kemp, Veerin R Sirihorachai, Stephanie The, Lawrence Delrosario, Jenny Lazarus, El-ad David Amir, Valerie Gunchick, Carlos Espinoza, et al. Multimodal mapping of the tumor and peripheral blood immune landscape in human pancreatic cancer. Nature Cancer, 1(11):1097–1112, 2020.

[33] Christopher J Halbrook, Marina Pasca di Magliano, and Costas A Lyssiotis. Tumor cross-talk networks promote growth and support immune evasion in pancreatic cancer. American Journal of Physiology-Gastrointestinal and Liver Physiology, 315(1):G27– G35, 2018.

