## Supplementary document for "SPACE: Spatially variable gene clustering adjusting for cell type effect for improved spatial domain detection"

**Contents**

**1 Score test for SVG detection** **3**

**2 SPACE algorithm** **4**

**3 Score statistic and its null distribution** **6**

**4 Multiplicity correction** **7**

**5 Analysis of human breast cancer dataset** **7**

### List of Figures

S1 The representative spatial genes in the human breast cancer dataset from SVG clusters detected by SPACE. . . . . . . . . . . . . . . . . . . . . . . . 8

S2 Steps to create a synthetic dataset from an annotated original Spatial dataset. We begin with a filtered gene expression count matrix and a location matrix where spots are annotated. The tissue region contains $N$ spots where gene expression measurements are measured for each of the $m$ genes. **Step 1:** Randomly select $m_1$ genes from the original dataset to be converted into new SVGs in the synthetic dataset. The remaining $m - m_1$ genes will serve as noise genes with no spatial pattern. **Step 2:** For each of the $m_1$ genes in the original dataset, sort and arrange the gene expression count values according to the reference domain structure. For the remaining genes, sort and arrange the count values randomly in the tissue region. . . . . . . . . . . . . . . . . . . . . . . . . . . . . . . . . 9

S3 The t-SNE plots[1] illustrate the separation of domain-specific SVGs by SPACE across 10 synthetic datasets. . . . . . . . . . . . . . . . . . . . . . 10

S4 The performance comparison between A)nnSVG[2] and B)SPACE is conducted for the simulation setup outlined in section 3.1.1 of the main manuscript. The gene groups are: Independent: Uncorrelated Gene group for genes without any spatial pattern(g1-g10), Correlated: Correlated Gene group for genes without any spatial pattern(g11-g20). Pattern 1-3: Correlated Gene group for genes with spatial pattern 1-3 (g21-g30,g31-g40,g41-g50). Pattern 4-6: Single gene with spatial pattern 4-6(g51-g53) The spatial pattern strength intensifies within each spatial gene group(pattern1-pattern3) as indicated by the triangles between the plots. The empirical power of the SVG detection step is evaluated for simulated datasets with AR(1) (Left) and compound symmetry (CS) (Right) correlation structures within the gene groups. . . . . . . . . . . . . . . . . . . . . . . . . . . . . . . . . 11

S5 The SVG-clusters detected from the DLPFC data using SPACE distinctly highlight the disparity between genes in the two main clusters. The representative genes in the second cluster demonstrate overexpression in the white matter region, while those in the first cluster exhibit overexpression in the other six cortex layers. Additionally, the three unique genes in the final row display a slightly different pattern compared to the genes in the main clusters. . . . . . . . . . . . . . . . . . . . . . . . . . . . . . 12

S6 Annotated and predicted spatial domains for all 12 DLPFC samples using SPACE, alongside ARI scores for each sample indicating prediction accuracy. 13
S7 t-SNE plots illustrating genes from all 12 DLPFC samples, with colors representing SVG cluster labels detected by SPACE. Across the majority of samples, distinct clustering is observed, indicating accurate separation of genes within different clusters. . . . . . . . . . . . . . . . . . . . . . . . . 14
S8 Analysis of PDAC Data using SPACE unveils three primary SVG clusters. Representative genes from each cluster (Cluster 1, Cluster 2, and Cluster 3) are showcased. The genes from these distinct clusters exhibit overexpression in three different regions. . . . . . . . . . . . . . . . . . . . 15
S9 Analysis of PDAC data: Pathway enrichment analysis of genes from A) list of all SVGs, B) list of top 3000 genes . . . . . . . . . . . . . . . . . . . 16

### 1 Score test for SVG detection

The Gaussian process (GP) regression model which models the normalized gene expression $y$ for a given gene using the following multivariate normal model:

$$p(y|\beta, \sigma_s^2, \delta, K) \sim N(y|X\beta, \sigma_s^2 K + \delta I), \tag{1}$$

where the covariance term is decomposed into a spatial and a non-spatial part, where $\delta I$ represents the non-spatial part and $\sigma_s^2 K$ is the spatial covariance matrix, whose $(i, j)^{th}$ element in the kernel matrix $K$ denotes the spatial similarity between the $i$th and $j$th spot calculated based on the corresponding coordinates $s_i$ and $s_j$. The choice of the kernel function plays a very important role in detecting the spatial correlation present in the gene expressions. $X^{N\times k}$ represents the covariate matrix, while $\beta^{k\times 1}$ denotes the array of corresponding coefficients.

As we mentioned in the main text, testing if a gene is a SVG is equivalent to testing $H_0 : \sigma_s^2 = 0$. The null hypothesis $H_0 : \sigma_s^2$ =0 can be tested using the variance-component score test which is the locally most powerful test[3]. The variance-covariance score statistic is:

$$Q = (y - X\hat{\beta})^T K (y - X\hat{\beta})$$

where $\hat{\beta}$ is the MLEs under the null model. Under the null hypothesis, the score statistic $Q$ follows a mixture of chi-square distributions [4], which can be closely approximated with the computationally efficient Davies' method[5]. More details about the test is provided in the next subsection.

In this manuscript, we utilize part of the code provided along with the SKAT paper

[4] which uses the same score test for the purpose of rare-variant association testing in genetic data.

**Kernel functions** A kernel function is defined as a function $K : \mathcal{X} \times \mathcal{X} \to \mathbb{R}$, where the kernel matrix $K = (k_{i,i'})_{i,i'=1}^{n}$ is symmetric and positive semidefinite with $k_{i,i'} = k(s_i, s_{i'})$. In this setting, $k(s_i, s_{i'})$ is a measure of similarity between the $i$th and the $i'$th subject. There are a variety of kernel functions to choose from, and the most simple one is the Linear kernel. The other useful kernels are Polynomial kernel, the Gaussian kernel and the cosine kernel. The functional forms of these kernels are summarized below:

- Linear kernel: $K(s_i, s_{i'}) = s_i^T s_{i'}$
- Polynomial kernel: $K(s_i, s_{i'}) = (s_i^T s_{i'} + c)^d$, where $c$,$d$ are the free parameters.
- Gaussian kernel: $K(s_i, s_{i'}) = exp\{-\|s_i - s_{i'}\|^2/l\}$, where $\|s_i - s_{i'}\|^2 = \sum_{j=1}^{p}(s_{ij} - s_{i'j})^2$ is the Euclidean distance, $l$ is a length scale parameter.
- Cosine kernel: $K(s_i, s_{i'}) = cos(2\pi\frac{\|s_i - s_{i'}\|^2}{\phi})$, where $\phi$ is the periodicity parameter.

**Choices of kernel functions:** We must define the kernel function in order to proceed with the hypothesis testing. As it is unknown which kernel will be best for the test, we employ the score test to evaluate the null hypothesis across various kernel functions with distinct kernel parameters. Gaussian and cosine kernels are typically effective in capturing spatial patterns. Following the method outlined in the SPARK paper [6], we compute five different length scale parameter values for the Gaussian Kernel and five different periodic parameter values for the cosine kernel. We conduct the test across ten different kernels and aggregate the resulting p-values using the Cauchy combination rule[7].

### 2 SPACE algorithm

Our model is built upon the Gaussian process model. Thus, we require normalized count matrix data with $m$ rows (genes) and $N$ columns (spots). We also need the spatial location matrix $L$ with $N$ rows, 2 columns ($X$ and $Y$ coordinates of spots).
**Step 1:** Detect SVGs based on model (1) defined in the main text. Suppose there are $m_1$ SVGs and denote the SVG list as $S_y$ with $|S_y| = m_1$ where $|\cdot|$ denotes the cardinality of a set.
**Step 2:** Start with the subset matrix denoted by $M_{SVG}^{m_1 \times N}$.
Repeat for $j = 1, \cdots, m_1$:

1). For the $j$th gene in $S_y$, find genes correlated with it using methods such as (1) SIS[8], (2) marginal correlation test (+ve correlation), (3) Elastic net[9][10], (4) SIS+Enet, or other methods. Denote the correlated gene list as $S_j$.

2). If $|S_j| = 0$, then the $j$th gene has an unique spatial pattern.

If $|S_j| <= 3$, then fit all genes in $S_j$ as the covariates in model (2) in the main text.

If $|S_j| > 3$, then get the $k_j$ PCs of genes in $S_j$ and fit them as covariates in model (2) in the main text.

3). Using the model (2) in the main text, conduct a score test to compute the p-value under 10 different kernels following the SPARK idea, then integrate these 10 p-values using the Cauchy combination rule[7] to get the final p-value. The output includes: a) For each SVG $j$, a list of correlated genes in $S_j$; and b) A list of unique SVGs as defined in 2.

**Step 3:**

1). Based on the output in step 2, a weighted graph structure is created where each SVG is a node. Each SVG $j$ in the output list in 3.a) has a common edge with all its dependent genes specified in list $S_j$ in the graph.

2). Clusters are determined from the weighted graph structure in 1) using the Leiden community detection algorithm[11].

3). The unique genes in output list 3.b) not connected with other nodes in the graph structure are allocated to the singleton set.

**SVG clustering with community detection algorithm:**

In many complex networks, nodes cluster and form relatively dense groups—often called communities, where the nodes within each community are more densely connected to each other than to the rest of the network. Such a modular structure is usually not known beforehand, making the detection of communities a challenging problem. One of the best-known methods for community detection is called modularity[12]. This method tries to maximize the difference between the actual number of edges in a community and the expected number of such edges. Community detection algorithms use various methodologies to partition networks into meaningful clusters, revealing insights into the relationships and interactions within the network. Different community detection algorithms like Louvain and Leiden are widely used in single-cell RNA sequencing (scRNA-seq) analysis for clustering single cells, aiding in the identification of distinct cell types and states.

The modularity is defined by:

$$\mathcal{H} = \frac{1}{2m} \sum_{c} (e_c - \gamma \frac{K_c^2}{2m})$$

where $m$ is the total number of edges in the network, $e_c$ is the actual number of edges in community $c$, the expected number of edges can be expressed as $\frac{K_c^2}{2m}$, where $K_c$ is the sum of the degrees of the nodes in community $c$. $\gamma > 0$ is a resolution parameter. Higher resolutions lead to more communities, while lower resolutions lead to fewer communities.

The Louvain method[13] is one of the most popular community detection algorithms due to its simplicity and effectiveness, typically operating based on modularity optimization. However, in recent years, some drawbacks of the Louvain algorithm have been identified[11], leading to the increased popularity of the Leiden algorithm. The Leiden method is more robust and accurate, making it better suited for analyzing complex networks where high-quality community detection is crucial.

In step 3 of the SPACE algorithm, we used the Leiden algorithm[11], implemented through the "igraph" R package[14],[15] with the default resolution parameter value $\gamma = 1$.

### 3 Score statistic and its null distribution

The Gaussian process model presented in Model (1) has many unknown parameters, for example $\delta$, the kernel parameter $\rho$, $\sigma_s^2$. The unknown parameters are estimated simultaneously by treating them as variance components in the linear mixed model and estimating them using the Restricted Maximum Likelihood(REML) model. The REML model was used to reduce bias in the variance components model [16][17][18].

The restricted log-likelihood[19][20] for the lmm in (1) is as follows:

$$L_R(\sigma_s^2, \delta, \rho) = -\frac{1}{2}log|\Sigma| - \frac{1}{2}log|X^T\Sigma^{-1}X| - \frac{1}{2}(y - X\beta)^T\Sigma^{-1}(y - X\beta) \quad (2)$$

where $\Sigma = \sigma_s^2 K + \delta I$.

The score statistic can be written in the form of $\tilde{Q}(\hat{\beta}, \hat{\delta}) - tr(HK)$[20], where $H = I - X(X^TX)^{-1}X^T$, $\hat{\beta}$ and $\hat{\delta}$ are the MLEs under the null model and $\tilde{Q}(\beta, \delta) = \frac{1}{2\delta}(y - X\beta)^T K(y - X\beta)$

Under $H_0 : \sigma_s^2 = 0$, the score statistic Q reduces to $y^T H^T KHy$ and

$$Q \sim \sum_{i=1}^{N} \lambda_i \chi_{i,1}^2 \quad (3)$$

where $\chi_{i,1}^2$ are independent $\chi_1^2$ random variables and $\lambda_1, \lambda_2, \cdots, \lambda_N$ are the eigenvalues of $\hat{\delta} H^{1/2} K H^{1/2}$.

The form of the null distribution of the score statistic in (3) follows from this argument [21]. Under $H_0$, $y \sim N(X\beta, \delta I)$, the covariate effects can be removed by projection, i.e., $\tilde{y} = Hy$. Now $\tilde{y} \sim N(0, \delta HH^T)$ and $K^{1/2}\tilde{y} \sim N(0, \delta K^{1/2}HH^TK^{1/2})$. Let $U$ be the matrix whose columns are the eigenvectors of $\hat{\delta}K^{1/2}HH^TK^{1/2}$ and the corresponding eigenvalues are $\tilde{\lambda}_1, \cdots, \tilde{\lambda}_N$. Therefore, $U^TK^{1/2}Hy \sim N(0, diag(\tilde{\lambda}_i))$ and $Q = (U^TK^{1/2}Hy)^T(U^TK^{1/2}Hy) \sim \sum_{i=1}^{N} \tilde{\lambda}_i\chi^2_{i,1}$. Using the fact that the eigenvalues of any matrix $A, AA^T, A^TA$ are the same and $H^2 = H$, it can be shown that $\tilde{\lambda}_i = \lambda_i$, the eigen values of $\hat{\delta}H^{1/2}KH^{1/2}$.

### 4 Multiplicity correction

At each stage of SPACE, our model is applied to each of the $m$ genes. To control the false discovery rate (FDR), multiplicity correction is required. We employ the Benjamini–Yekutieli (BY) procedure, known for its effectiveness under arbitrary correlation conditions, to obtain adjusted p-values to claim significant genes [22].

### 5 Analysis of human breast cancer dataset

We analyzed another spatial transcriptomics dataset of human breast cancer. We obtained the data from the SPARK[6] GitHub repository at https://github.com/xzhoulab/SPARK-Analysis (specifically the 'Layer2_BC_count_matrix-1.tsv' file). The dataset can also be downloaded from Spatial Transcriptomics Research (http://www.spatialtranscriptomicsresearch.org). This dataset contains 14,789 genes measured across 251 spots. Our preprocessing involved filtering out genes with expression levels below 10% across the array spots and retaining spots with a total read count > 10. Following these criteria, our analysis focused on a final set of 5,262 genes observed across 250 spots within the breast cancer dataset.

Using our framework SPACE, we identified 724 spatially relevant genes in the initial step. In comparison, SPARK[6] detected 290 genes, SPARK-G detected 244 genes, nnSVG[2] detected 592 genes, and SPARK-X[23] detected 901 genes. The 724 genes were subsequently classified into three main clusters(containing 369, 320, and 14 genes respectively) in the second step of SPACE framework, along with a few unique singleton genes. The representative genes from each cluster are visualized in Figure S1. Genes in cluster 1 exhibited higher expression levels in the lower tissue region, while cluster 2 genes were overexpressed in the middle tissue region, and cluster 3 genes highlighted the upper tissue region. Additionally, the unique genes showcased distinct expression patterns in the third row. This figure confirms the superior performance of our SPACE method.

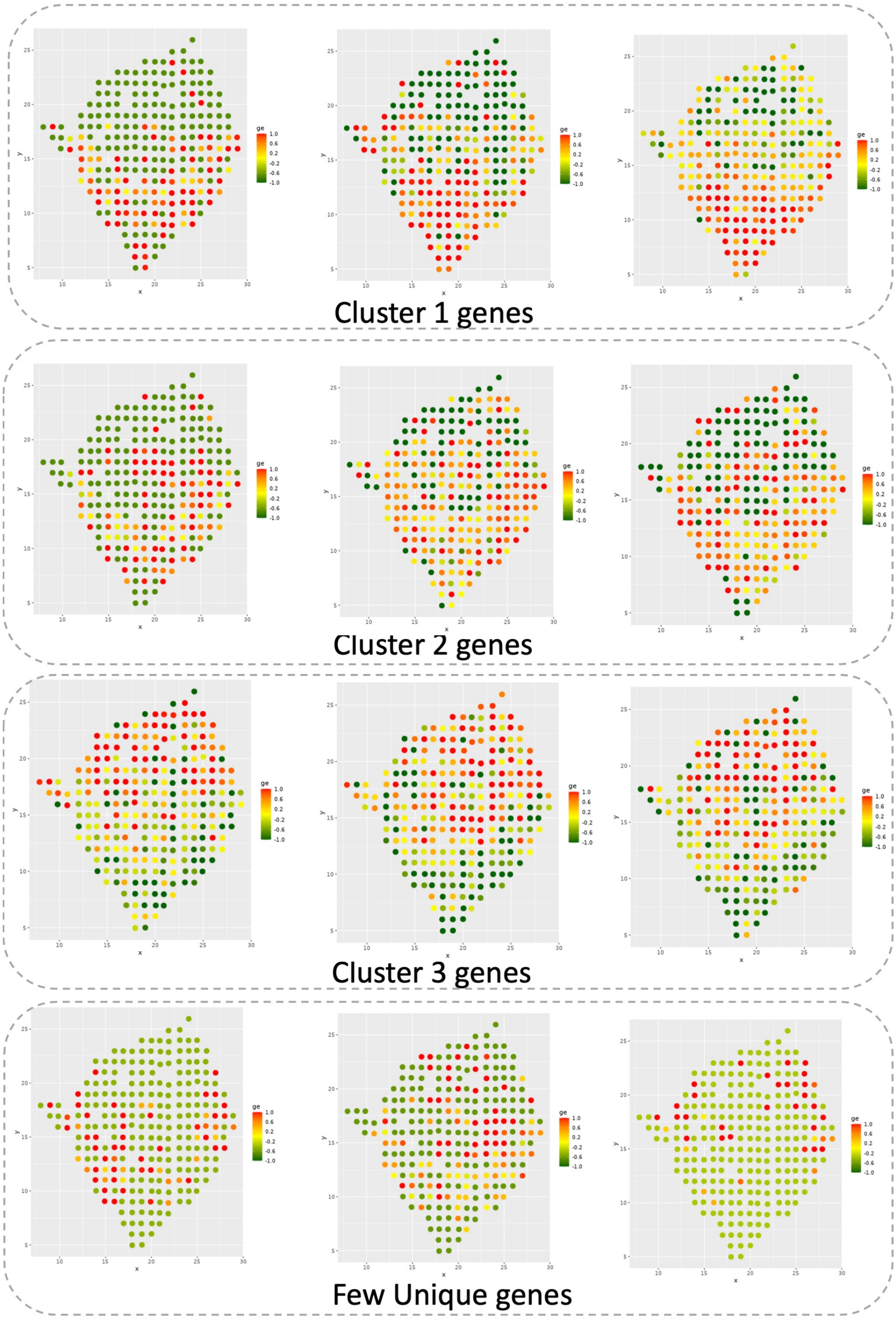


Figure S1: The representative spatial genes in the human breast cancer dataset from SVG clusters detected by SPACE.

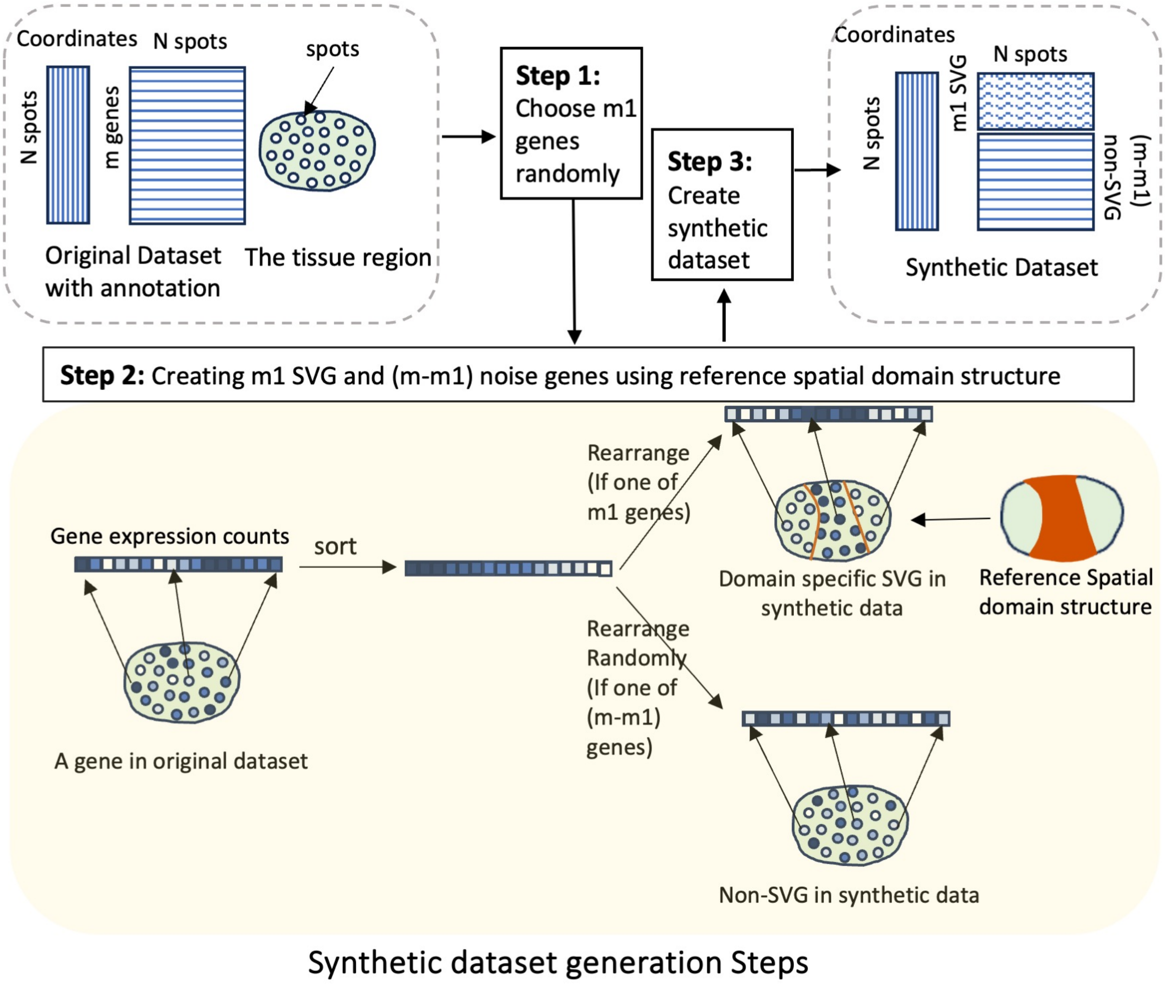


Figure S2: Steps to create a synthetic dataset from an annotated original Spatial dataset. We begin with a filtered gene expression count matrix and a location matrix where spots are annotated. The tissue region contains $N$ spots where gene expression measurements are measured for each of the $m$ genes. **Step 1:** Randomly select $m_1$ genes from the original dataset to be converted into new SVGs in the synthetic dataset. The remaining $m - m_1$ genes will serve as noise genes with no spatial pattern. **Step 2:** For each of the $m_1$ genes in the original dataset, sort and arrange the gene expression count values according to the reference domain structure. For the remaining genes, sort and arrange the count values randomly in the tissue region.

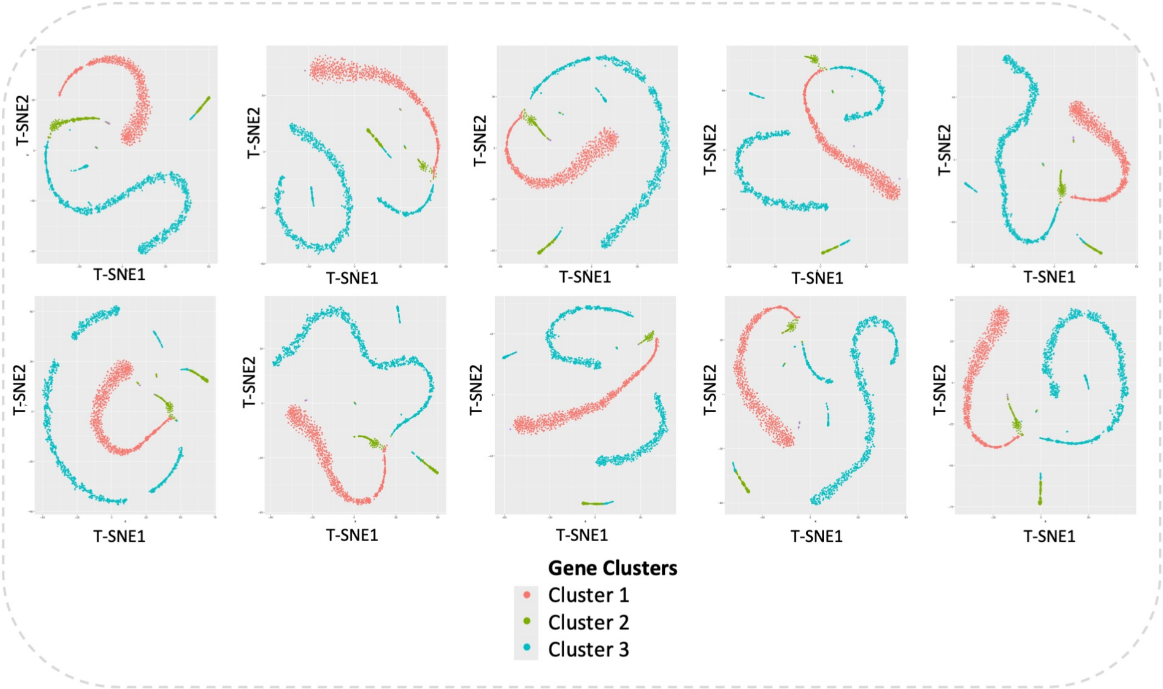


Figure S3: The t-SNE plots[1] illustrate the separation of domain-specific SVGs by SPACE across 10 synthetic datasets.

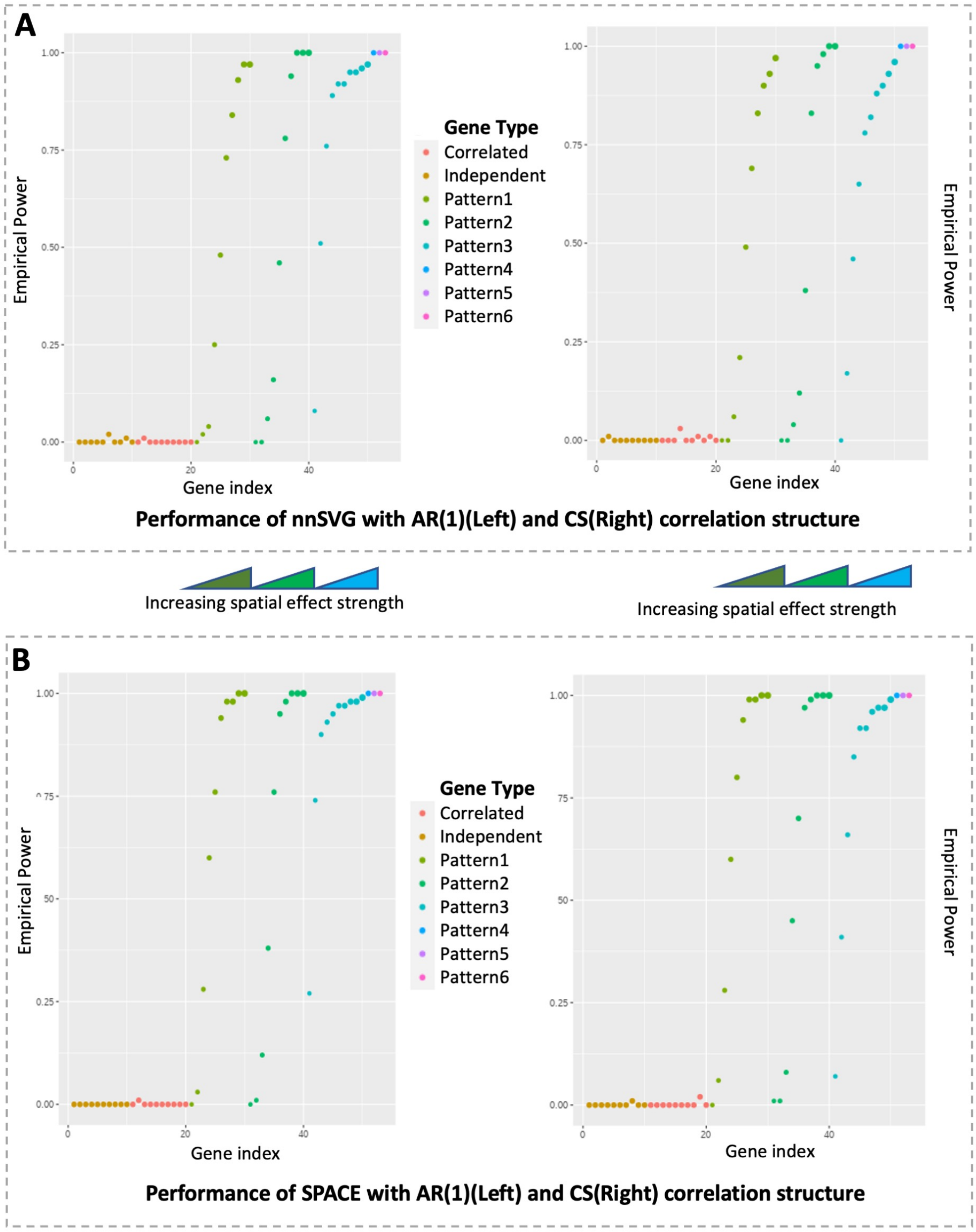


Figure S4: The performance comparison between A)nnSVG[2] and B)SPACE is conducted for the simulation setup outlined in section 3.1.1 of the main manuscript. The gene groups are: Independent: Uncorrelated Gene group for genes without any spatial pattern(g1-g10), Correlated: Correlated Gene group for genes without any spatial pattern(g11-g20). Pattern 1-3: Correlated Gene group for genes with spatial pattern 1-3 (g21-g30,g31-g40,g41-g50). Pattern 4-6: Single gene with spatial pattern 4-6(g51-g53) The spatial pattern strength intensifies within each spatial gene group(pattern1-pattern3) as indicated by the triangles between the plots. The empirical power of the SVG detection step is evaluated for simulated datasets with AR(1) (Left) and compound symmetry (CS) (Right) correlation structures within the gene groups.

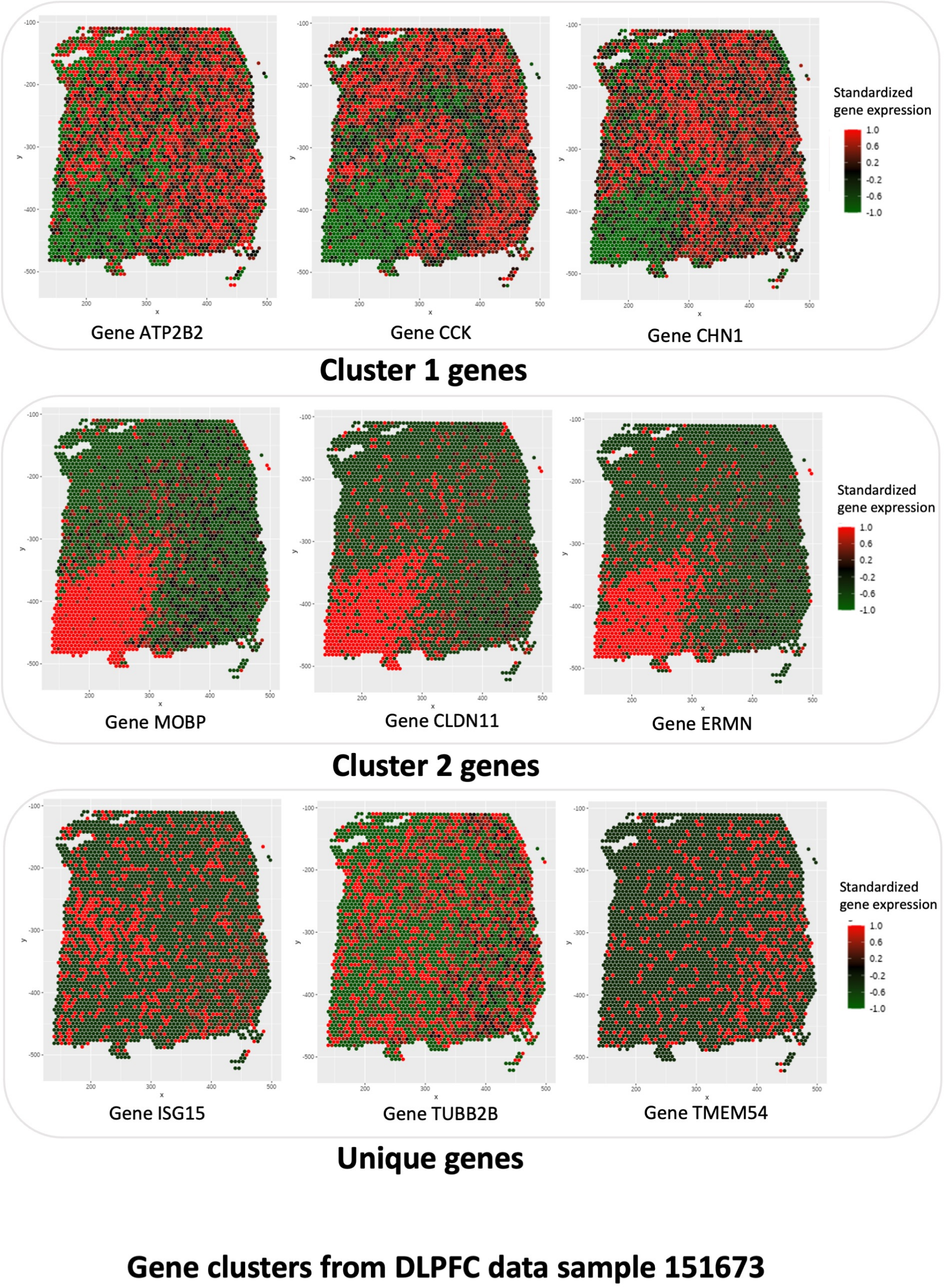


Figure S5: The SVG-clusters detected from the DLPFC data using SPACE distinctly highlight the disparity between genes in the two main clusters. The representative genes in the second cluster demonstrate overexpression in the white matter region, while those in the first cluster exhibit overexpression in the other six cortex layers. Additionally, the three unique genes in the final row display a slightly different pattern compared to the genes in the main clusters.

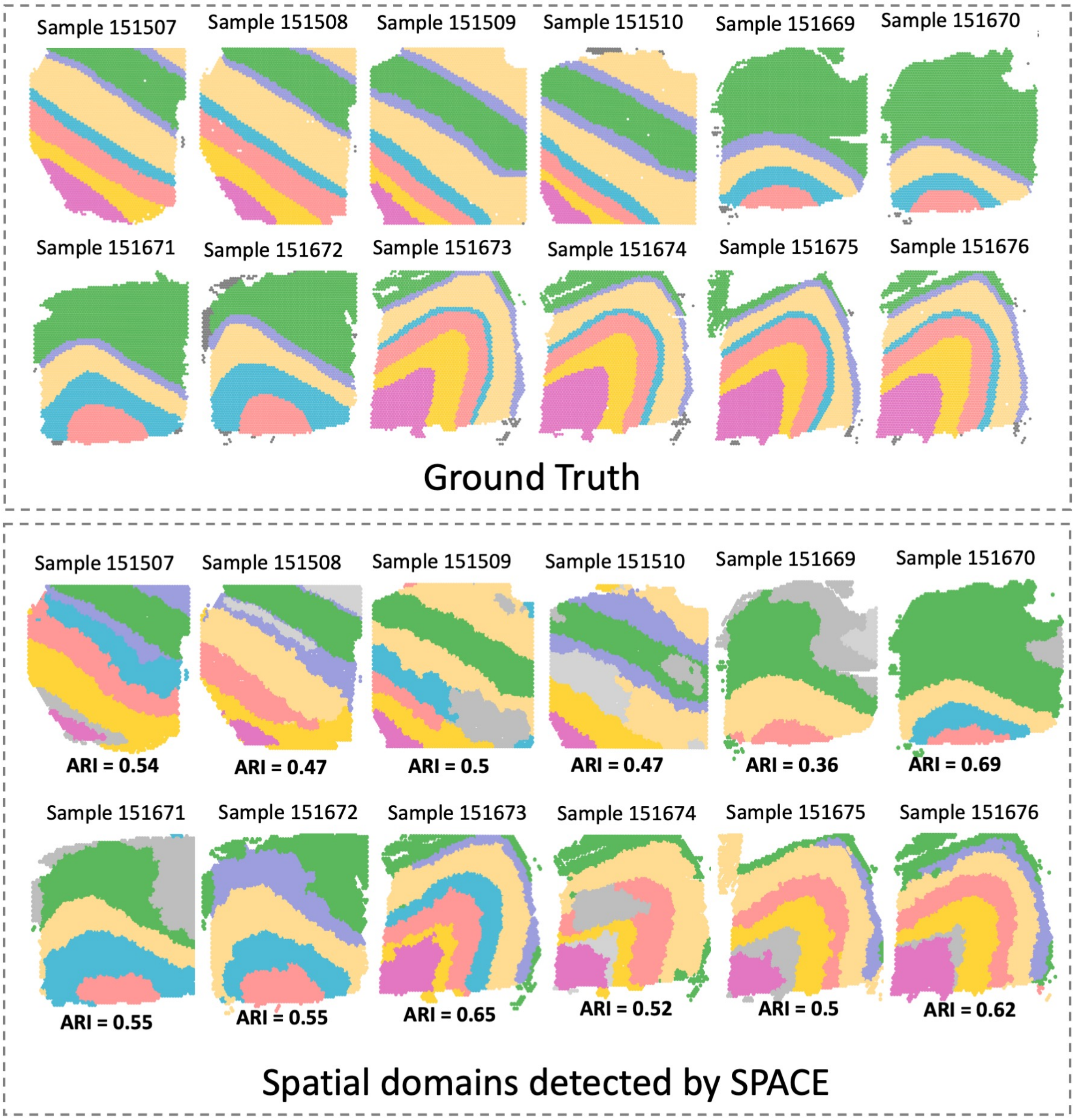


Figure S6: Annotated and predicted spatial domains for all 12 DLPFC samples using SPACE, alongside ARI scores for each sample indicating prediction accuracy.

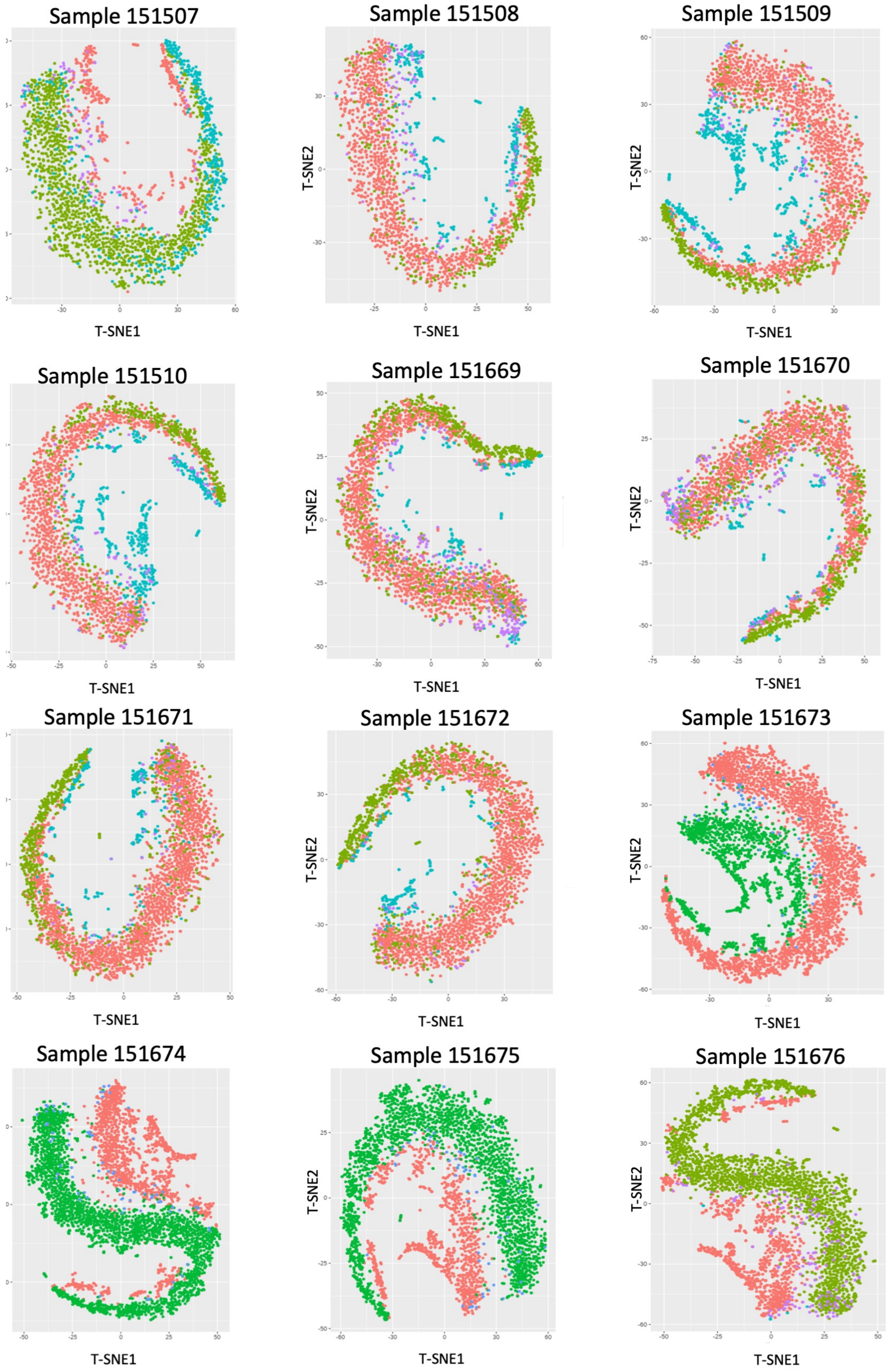


Figure S7: t-SNE plots illustrating genes from all 12 DLPFC samples, with colors representing SVG cluster labels detected by SPACE. Across the majority of samples, distinct clustering is observed, indicating accurate separation of genes within different clusters.

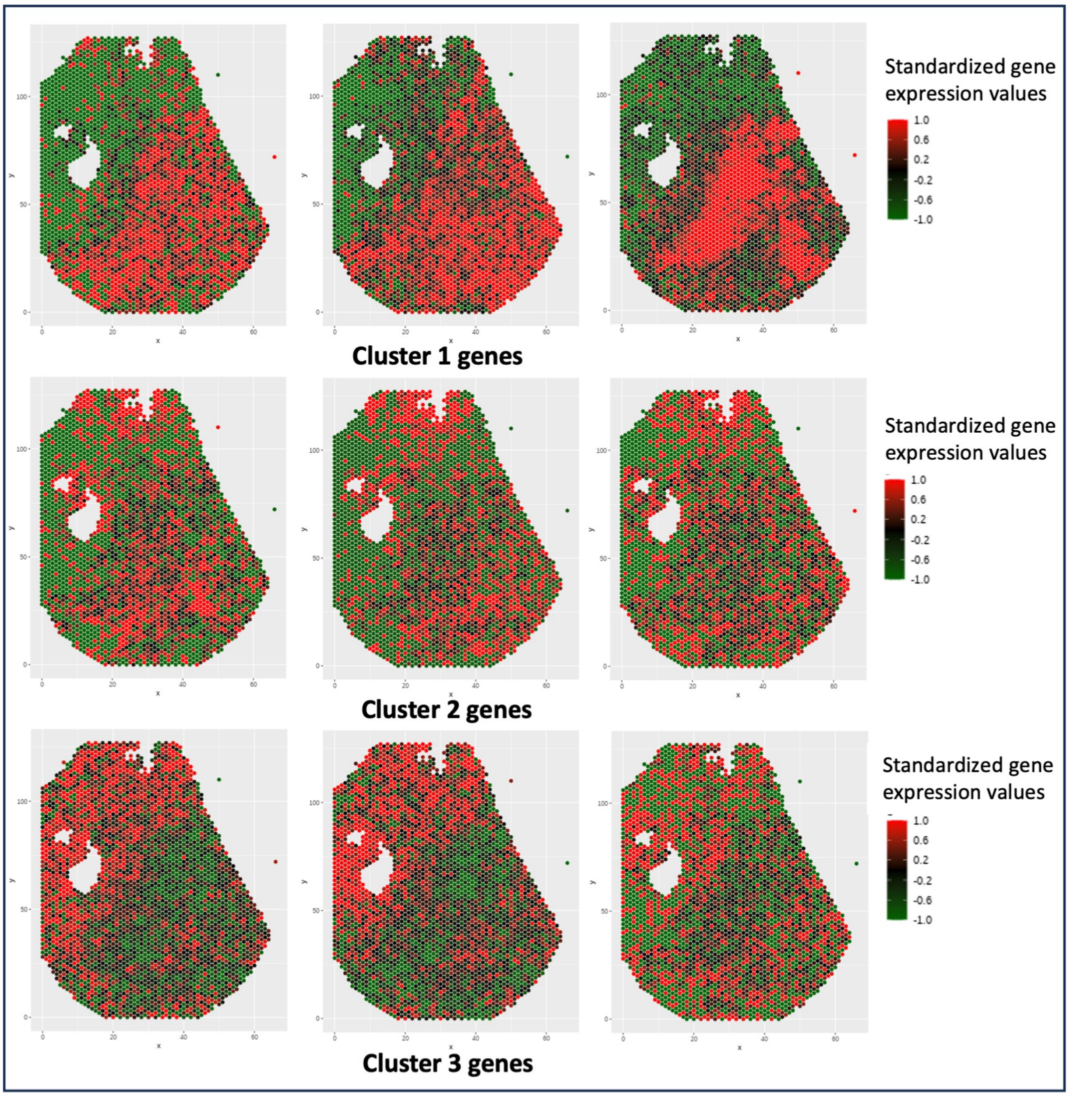


Figure S8: Analysis of PDAC Data using SPACE unveils three primary SVG clusters. Representative genes from each cluster (Cluster 1, Cluster 2, and Cluster 3) are showcased. The genes from these distinct clusters exhibit overexpression in three different regions.

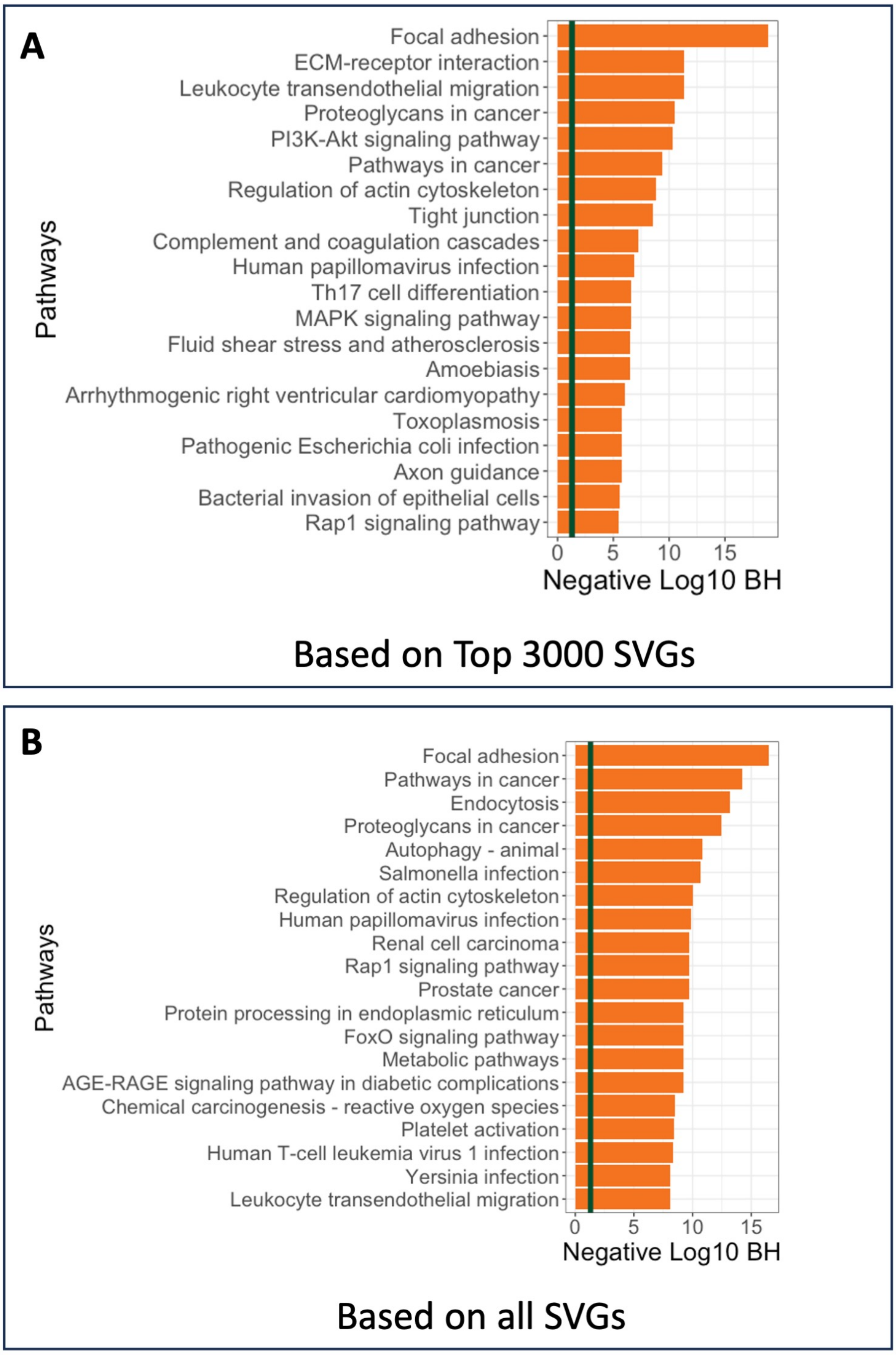


Figure S9: Analysis of PDAC data: Pathway enrichment analysis of genes from A) list of all SVGs, B) list of top 3000 genes

### References


[1] Laurens Van der Maaten and Geoffrey Hinton. Visualizing data using t-sne. *Journal of machine learning research*, 9(11), 2008.

[2] Lukas M Weber, Arkajyoti Saha, Abhirup Datta, Kasper D Hansen, and Stephanie C Hicks. nnsvg for the scalable identification of spatially variable genes using nearest-neighbor gaussian processes. *Nature communications*, 14(1):4059, 2023.

[3] Xihong Lin. Variance component testing in generalised linear models with random effects. *Biometrika*, 84(2):309–326, 1997.

[4] Michael C Wu, Seunggeun Lee, Tianxi Cai, Yun Li, Michael Boehnke, and Xihong Lin. Rare-variant association testing for sequencing data with the sequence kernel association test. *The American Journal of Human Genetics*, 89(1):82–93, 2011.

[5] Robert B Davies. The distribution of a linear combination of $\chi 2$ random variables. *Journal of the Royal Statistical Society Series C: Applied Statistics*, 29(3):323–333, 1980.

[6] Shiquan Sun, Jiaqiang Zhu, and Xiang Zhou. Statistical analysis of spatial expression patterns for spatially resolved transcriptomic studies. *Nature methods*, 17(2):193–200, 2020.

[7] Yaowu Liu, Sixing Chen, Zilin Li, Alanna C Morrison, Eric Boerwinkle, and Xihong Lin. Acat: a fast and powerful p value combination method for rare-variant analysis in sequencing studies. *The American Journal of Human Genetics*, 104(3):410–421, 2019.

[8] Jianqing Fan and Jinchi Lv. Sure independence screening for ultrahigh dimensional feature space. *Journal of the Royal Statistical Society Series B: Statistical Methodology*, 70(5):849–911, 2008.

[9] Jerome Friedman, Trevor Hastie, and Rob Tibshirani. Regularization paths for generalized linear models via coordinate descent. *Journal of statistical software*, 33(1):1, 2010.

[10] Noah Simon, Jerome Friedman, Trevor Hastie, and Rob Tibshirani. Regularization paths for cox's proportional hazards model via coordinate descent. *Journal of statistical software*, 39(5):1, 2011.

[11] Vincent A Traag, Ludo Waltman, and Nees Jan Van Eck. From louvain to leiden: guaranteeing well-connected communities. *Scientific reports*, 9(1):5233, 2019.

[12] Mark EJ Newman and Michelle Girvan. Finding and evaluating community structure in networks. *Physical review E*, 69(2):026113, 2004.

[13] Vincent D Blondel, Jean-Loup Guillaume, Renaud Lambiotte, and Etienne Lefebvre. Fast unfolding of communities in large networks. *Journal of statistical mechanics: theory and experiment*, 2008(10):P10008, 2008.

[14] Gabor Csardi and Tamas Nepusz. The igraph software package for complex network research. *InterJournal*, Complex Systems:1695, 2006.

[15] Gábor Csárdi, Tamás Nepusz, Vincent Traag, Szabolcs Horvát, Fabio Zanini, Daniel Noom, and Kirill Müller. *igraph: Network Analysis and Visualization in R*, 2024. R package version 2.0.3.

[16] Eugene Demidenko. *Mixed models: theory and applications with R*. John Wiley & Sons, 2013.

[17] David A Harville. Bayesian inference for variance components using only error contrasts. *Biometrika*, 61(2):383–385, 1974.

[18] H Desmond Patterson and Robin Thompson. Recovery of inter-block information when block sizes are unequal. *Biometrika*, 58(3):545–554, 1971.

[19] Nan M Laird and James H Ware. Random-effects models for longitudinal data. *Biometrics*, pages 963–974, 1982.

[20] Dawei Liu, Xihong Lin, and Debashis Ghosh. Semiparametric regression of multidimensional genetic pathway data: least-squares kernel machines and linear mixed models. *Biometrics*, 63(4):1079–1088, 2007.

[21] Regev Schweiger, Omer Weissbrod, Elior Rahmani, Martina Müller-Nurasyid, Sonja Kunze, Christian Gieger, Melanie Waldenberger, Saharon Rosset, and Eran Halperin. Rl-skat: an exact and efficient score test for heritability and set tests. *Genetics*, 207(4):1275–1283, 2017.

[22] Yoav Benjamini and Daniel Yekutieli. The control of the false discovery rate in multiple testing under dependency. *Annals of statistics*, pages 1165–1188, 2001.

[23] Jiaqiang Zhu, Shiquan Sun, and Xiang Zhou. Spark-x: non-parametric modeling enables scalable and robust detection of spatial expression patterns for large spatial transcriptomic studies. *Genome biology*, 22(1):184, 2021.
